# Maternally-regulated gastrulation as a source of variation contributing to cavefish forebrain evolution

**DOI:** 10.1101/410563

**Authors:** Jorge Torres-Paz, Julien Leclercq, Sylvie Rétaux

**Affiliations:** Paris-Saclay Institute of Neuroscience, CNRS UMR9197, Université Paris-Sud and Université Paris-Saclay, Gif-sur-Yvette, France

**Keywords:** *Astyanax mexicanus*, developmental evolution, heterochrony, *dkk1b*, Wnt signaling, prechordal plate, notochord, maternal transcriptome

## Abstract

Sequential developmental events, starting from the moment of fertilization, are crucial for the acquisition of animal body plan. Subtle modifications in such early events are likely to have a major impact in later morphogenesis, bringing along morphological diversification. Here, comparing the blind cave and the surface morphotypes of *Astyanax mexicanus* fish, we found heterochronies during gastrulation, producing organizer and axial mesoderm tissues with different properties, including differences in expression of *dkk1b*, that may have contributed to cavefish brain evolution. These variations observed during gastrulation depend fully on maternal factors, whereas later phenotypic differences in neural development became progressively hidden when zygotic genes take the control over development. Transcriptomic analysis of fertilized eggs from both morphotypes and reciprocal F1 hybrids showed a strong and specific maternal signature. Our work strongly suggests that maternal effect genes and developmental heterochronies occurring during gastrulation have impacted morphological brain change during cavefish evolution.

## Introduction

Gastrulation is a fundamental process in organism development, leading to the establishment of the embryonic germ layers (endoderm, mesoderm and ectoderm) and the basic organization of the body plan. Although in vertebrates early embryonic development has adopted highly diverse configurations, gastrulation proceeds through evolutionary conserved morphogenetic movements, including the spreading of blastoderm cells (epiboly), the internalization of mesoderm and endoderm, convergent movements towards the prospective dorsal side and extension along the antero-posterior axis (convergence and extension, respectively) (Solnica-Krezel, 2005). Internalization of mesendodermal cells takes place through the blastopore, structurally circumferential in anamniotes (fishes and amphibians) and lineal in avian and mammalian amniotes (primitive streak).

A critical step for gastrulation to proceed is the establishment of the embryonic organizer (Spemann-Mangold organizer in frogs, shield in fishes, Hensen’s node in birds and node in mammals), a signaling center essential to instruct the formation of the body axis. In fishes and amphibians the induction of the embryonic organizer in the prospective dorsal side occurs downstream to earlier developmental events, driven by maternal determinants deposited in the oocyte during maturation in the ovaries (Kelly et al., 2000; Nojima et al., 2004; Zhang et al., 1998). From the organizer will emerge the axial mesoderm, a structure that spans the complete rostro-caudal extent of the embryo, with the prechordal plate anteriorly and the notochord posteriorly. The axial mesoderm is the signaling center that will induce vertically the neural plate/tube in the overlying ectoderm.

The prechordal plate is key for the patterning of the forebrain, through the regulated secretion of morphogens including sonic hedgehog (shh), Fibroblast growth factors (fgf), and inhibitors of the Wingless-Int (Wnt) pathway like dickkopf1b (dkk1b) and secreted frizzled-related proteins (sFRP). Along its rostral migration, the prechordal plate is required for sequential patterning of forebrain elements (García-Calero et al., 2008; Puelles & Rubenstein, 2015), demonstrating a temporal and spatial requirement of this migratory cell population for brain development from gastrulation onwards.

Within the central nervous system, the forebrain plays a key role in processing sensory information coming from the environment and controlling higher cognitive functions. During evolution and across species, different forebrain modules have experienced impressive morphological modifications according to ecological needs, however the basic *Bauplan* to build the forebrain has been conserved. Temporal (heterochronic) and spatial (heterotopic) variation in the expression of regionalization genes and morphogens during embryogenesis have sculpted brain shapes along phylogeny (Bielen et al., Houart, 2017; Rétaux et al., 2013).

An emergent model organism to study the impact of early embryogenesis on brain evolution at the microevolutionary scale is the characid fish *Astyanax mexicanus*. This species exists in two different eco-morphotypes distributed in Central and North America: a “wild type” river-dwelling fish (surface fish) and several geographically-isolated troglomorphic populations (cavefish), living in total and permanent darkness (Mitchell et al., 1977; Elliott, 2018). Fish from the cave morphotype can be easily identified because they lack eyes and pigmentation. As a result of the absence of visual information the cavefish has evolved mechanisms of sensory compensation, as enhanced chemosensory and mechanosensory sensibilities (Hinaux et al., 2016; Yoshizawa et al, 2010). Sensory and other behavioral adaptations may have allowed them to increase the chances of finding food and mates in caves. Such behavioral changes are associated with morphological modifications such as larger olfactory sensory organs (Blin et al., 2018; Hinaux et al., 2016), increased number of facial mechanosensory neuromasts (Yoshizawa et al., 2014) and taste buds (Varatharasan et al., 2009), and modified serotonergic and orexinergic neurotransmission systems (Alié et al., 2018; Elipot et al., 2014; Jaggard et al., 2018). Remarkably, such morphological and behavioral adaptations have a developmental origin, mainly due to heterotopic and heterochronic differences in the expression of signaling molecules from midline organizers at the end of gastrulation, at the “neural plate” or bud stage. Subtle differences in *shh* and *fgf8* expression domains, larger and earlier respectively in cavefish compared to surface fish, affect downstream processes of gene expression, morphogenetic movements during neurulation and cell differentiation, driving the developmental evolution of cavefish nervous system (Hinaux et al., 2016; Menuet et al., 2007; Pottin et al., 2011; Ren et al., 2018; Yamamoto et al, 2004). As these differences in genes expressed in the midline are already manifest in embryos at the end of body axis formation, we postulated that they should stem from earlier developmental events during axis formation and gastrulation.

In order to search for variations in precocious ontogenetic programs leading to phenotypic evolution observed in *A. mexicanus* morphotypes, here we performed a systematic comparison of the gastrulation process in cave and surface embryos. We found that in the cavefish, migration of different mesodermal cell populations is more precocious, prompting us to go further backwards in embryogenesis and to investigate maternal components. Taking advantage of the inter-fertility of the two morphotypes we compared gastrulation, forebrain phenotypes and maternal transcriptomes in embryos obtained from reciprocal crosses between cavefish/surface fish males/females. We found that maternal factors present in the egg contribute greatly to the evolution of cavefish gastrulation and subsequent forebrain developmental evolution.

## Results

### Molecular identity of the gastrula margin in A. mexicanus

In the zebrafish the embryonic organizer/shield becomes morphologically evident at the prospective dorsal margin of the blastopore right after the epiboly has covered half of the yolk cell (50% epiboly), a stage that coincides with the initiation of the internalization of mesendodermal precursors. We studied the expression of genes involved in the establishment of the organizer in the two *A. mexicanus* morphotypes at the equivalent stage by ISH, in order to search for early differences.

First, at 50% epiboly, the inhibitor of the Wnt signaling pathway, *Dkk1b*, was expressed in a strikingly different pattern in the two morphs. In the surface fish, *dkk1b* expression was observed at the dorsal margin in two groups of cells separated by a gap in the center (Figure 1A), a pattern observed in the majority of the embryos (around 70%; Figure 1C blue). In the cavefish, a single central spot of variable extension (Figure 1B) was observed in most of the samples analyzed (around 70%; Figure 1C red). A minority of embryos of each morphotype showed an intermediate pattern corresponding to a line of positive cells without a clear interruption (not shown, Figure 1C green). To interpret this *dkk1b* pattern difference between the two morphs, fluorescent ISH and confocal imaging was performed. In cavefish at 50% epiboly, the *dkk1b*+ cells were already internalized under the dorsal aspect of the margin (Figure 1D-D’’ **and** 1E, E’), revealing a precocious internalization process as compared to surface fish.

**Figure 1.**
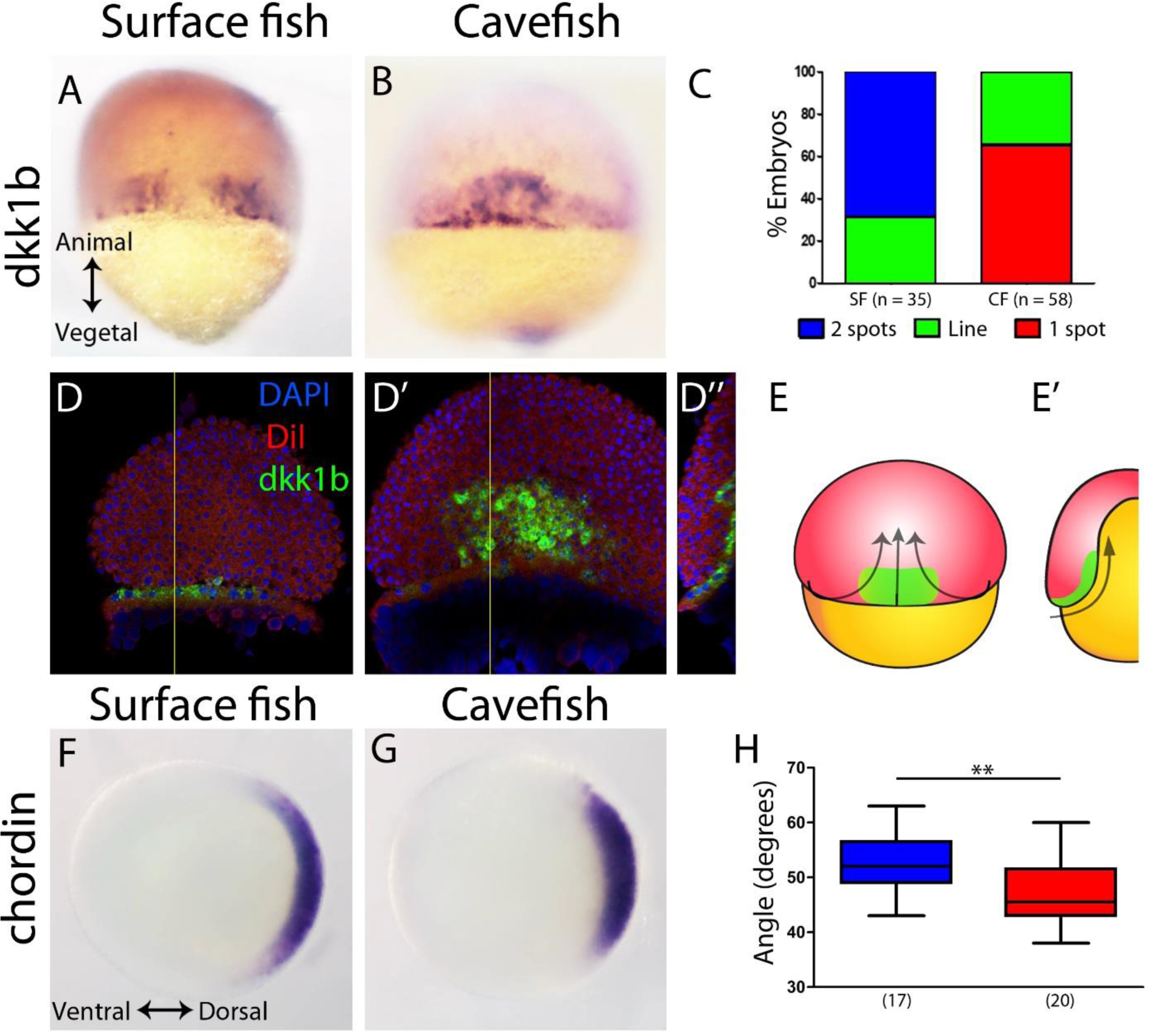
Expression of genes in the organizer at 50% epiboly in surface fish and cavefish. (A-B) Expression of *dkk1b* in surface fish (A) and cavefish (B) in dorsal view. (C) Quantification of the expression patterns observed in each morphotype. The Y-axis indicates the percentage of embryos belonging to each of the categories and the number of embryos analyzed is indicated. “Two spots” (blue) is the pattern observed in A, “1 spot” (red) is the pattern observed in B, and “Line” is an intermediate profile (not shown). (D-D’’) Confocal optical sections of superficial (D) and deep (D’) planes and orthogonal section (D’’) at the level of the yellow line in D-D’ of a cavefish embryo stained with DiI (red) and DAPI (blue) after fluorescent ISH to *dkk1b*. Representation of the cell movements of convergence and internalization (arrows) in a dorsal view (E) and in a section (E’), with the *dkk1b+* cells in green. (F-G) Expression of *chordin* in surface fish (F) and cavefish (G) in animal view. (H) Quantification of the angle covered *chordin* expression pattern. A, B, D, D’ are dorsal views, animal pole upwards. F, G are animal views, dorsal to the right. Mann-Whitney test was performed in H, ** = 0.0083.

Chordin is a dorsalizing factor, inhibitor of the Bmp pathway. In *A. mexicanus* it is expressed broadly in the dorsal side (Figure 1F, G), similarly to the pattern in zebrafish embryos (Langdon & Mullins, 2011; Miller-Bertoglio et al., 1997). In surface fish embryos *chordin* expression extended more ventrally than in cavefish (Figure 1F-H), as quantified by measuring the angle of expression in an animal view (Figure 1 - figure supplement 1). From a dorsal view *chordin* showed a slightly larger extension in the vegetal to animal axis, although not significant (not shown). This difference in *chordin* pattern extension suggested that convergence towards the dorsal pole was more advanced in cavefish.

Lefty1 is part of a feedback loop regulating nodal signaling activity, involved in axial mesoderm formation and lateral asymmetry establishment (Bisgrove et al., 1999; Meno et al., 1998). In *A. mexicanus* embryos *lefty1* expression was observed in the dorsal margin at 50% epiboly (Figure 1 - figure supplement 2A-B). The ventro-dorsal extension of *lefty1* expression was similarly variable in both morphotypes at this stage (not shown) and no significant differences were observed in the vegetal-animal extension (Figure 1 - figure supplement 2C).

We also compared the expression of 3 genes involved in notochord development: *floating head* (*flh*), *no-tail* (*ntl*) and *brachyury* (*bra*) (Glickman et al., 2003; Schulte-Merker et al., 1994; Talbot et al., 1995). At 50% epiboly the homeobox gene *flh* showed localized expression in the dorsal margin (Figure 1 - figure supplement 2D-E), without differences neither in width nor in height when compared between morphotypes (Figure 1 - figure supplement 2F). At the same stage *ntl* and *bra* expression extended homogenously all around the margin (blastopore), hindering the identification of the prospective dorsal side (Figure 1 - figure supplement 2G, H **and** I, J, respectively). No differences were observed between surface fish and cavefish.

In zebrafish, *dkk1b* is expressed in two spots in the embryonic organizer (Hashimoto et al., 2000) similarly to the surface fish condition (Figure 1A) - although the gap is less pronounced. We reasoned that the size of the “*dkk1b* gap” may vary due to differences in dorsal convergence and internalization of mesodermal lineages during gastrulation, relative to epiboly. The narrower domain of *chordin* expression observed in cavefish compared to surface fish also supported this hypothesis. To test this idea, we next analyzed the expression of axial mesodermal markers during subsequent stages of gastrulation.

### Mesoderm migration timing in A. mexicanus morphotypes

The EVL (enveloping layer) and YSL (yolk syncytial layer) drive epiboly movements that engulf the yolk cell (Bruce, 2016). Axial mesoderm precursors are mobilized from the dorsal organizer towards the rostral extreme of the embryo (animal pole), migrating in between the YSL and the epiblast (prospective neurectoderm). Since these events are important for the induction and patterning of the neural tube, we compared in detail the process of axial mesoderm migration in *A. mexicanus* morphotypes using markers of different mesodermal populations, always taking the percentage of epiboly as reference to stage embryos.

The *dkk1b* patterns in the two morphs were also clearly different towards mid-gastrulation. In surface fish, the two clusters observed at 50% epiboly began to coalesce at the midline at 70% epiboly (Figure 2A), whereas in cavefish*dkk1b* expressing cells became grouped dorsally and leading cells were more advanced towards the animal pole (arrow in Figure 2B). At 80% epiboly, *dkk1b*+ cells in the cavefish were close to their final position in the anterior prechordal plate at the rostral end of the embryonic axis (arrow Figure 2E). At the same stage, leading cells expressing *dkk1b* in the surface fish (arrow Figure 2D) had reached a similar distance as they did in cavefish at 70% epiboly (compare values in Figure 2F and **2C**). These expression profiles indicated that even though at 50% epiboly *dkk1b* expression appears very divergent in the two morphotypes, the cellular arrangement observed later on are similar, although always more advanced in the cavefish.

**Figure 2.**
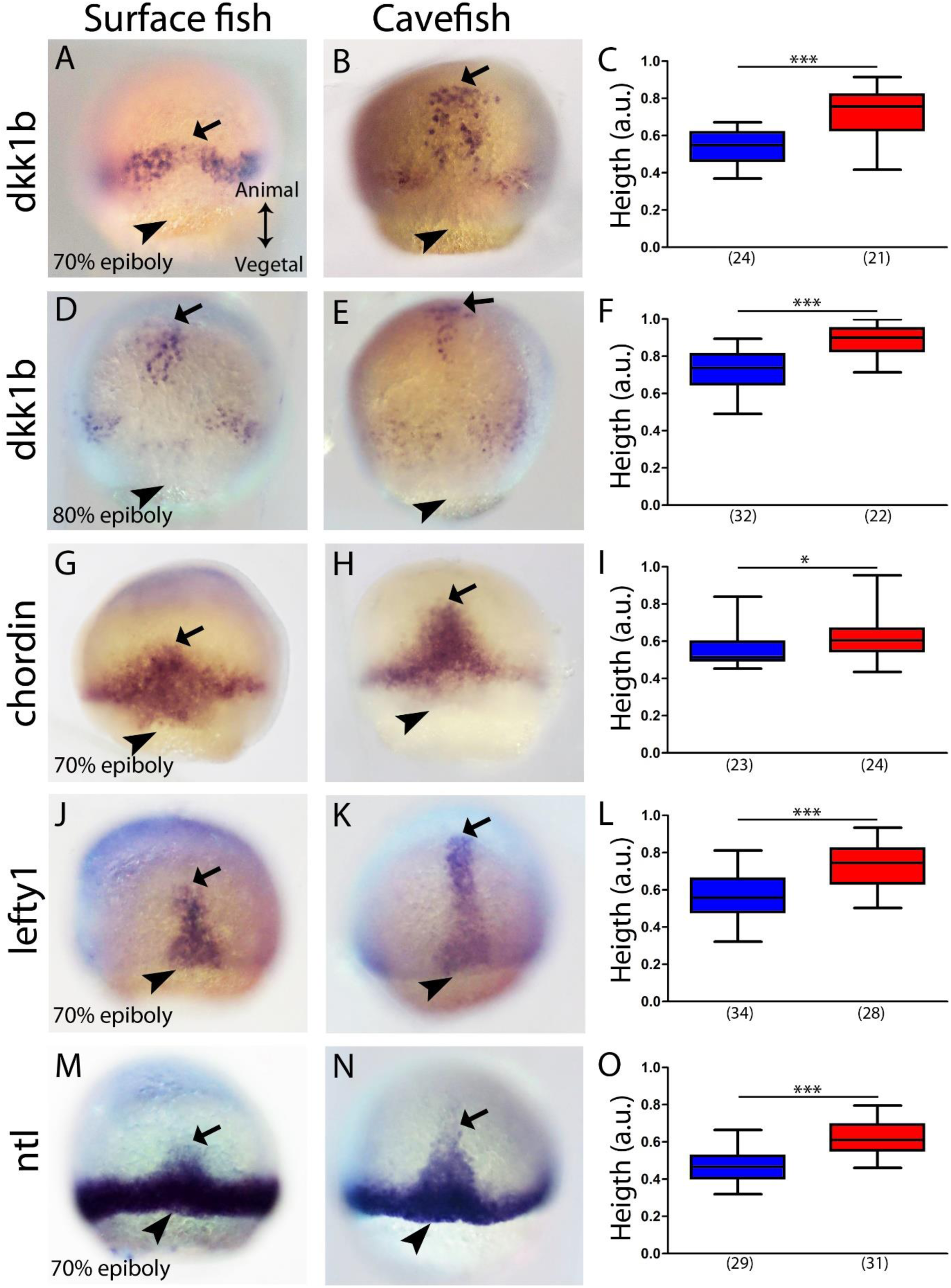
Expression of axial mesodermal genes during mid-gastrulation in surface fish and cavefish. (A-B, D-E) Expression of *dkk1b* in surface fish (A, D) and cavefish (B, E) at 70 and 80 % epiboly (A-B and D-E, respectively). (C, F) Quantification of height in *dkk1b* labeled embryos at 70 and 80% epiboly (C and F, respectively). (G-H) Expression of *chordin* in surface fish (G) and cavefish (H) at 70% epiboly. (I) Quantification of height in *chordin* labeled embryos at 70% epiboly. (J-K) Expression of *lefty1* in surface fish (J) and cavefish (K) at 70% epiboly. (L) Quantification of height in *lefty1* labeled embryos at 70% epiboly. (M-N) Expression of *ntl* in surface fish (M) and cavefish (N) at 70% epiboly. (O) Quantification of height in *ntl* labeled embryos at 70% epiboly. Embryos in dorsal views, animal pole upwards. Mann-Whitney test were performed. *** < 0.0001, * = 0.0167 (I).

The same analysis was performed at 70% epiboly for the markers *chordin* (Figure 2G-I), *lefty1* (Figure 2J-L) and *ntl* (Figure 2M-O). These 3 genes showed a greater height value of their expression domain in cavefish than in surface fish embryos. This further suggested that at equivalent stages during gastrulation anteroposterior axis formation is more advanced in cavefish.

Next, we wondered if the observed phenotype for the cavefish axial mesoderm also extends to the neighboring paraxial mesoderm, *i.e.*, the mesodermal tissue located laterally that will give rise to the somites (presomitic mesoderm). We analyzed the expression of *myoD* and *mesogenin 1* (*msgn1*), two genes coding for bHLH transcription factors required for early specification of myogenic tissue (Weinberg et al., 1996; Yabe and Takada, 2012). In *A. mexicanus*, at mid-gastrulation *myoD* was expressed in two domains, triangular in shape, on both sides of the dorsal axial mesoderm, corresponding to the central gap without expression (Figure 3A-D). The height value of the expression domain was higher in cavefish embryos both at 70% and 80% epiboly compared to the surface fish (Figure 3E), whereas the central/dorsal non-expressing zone was wider in the surface fish at both stages (Figure 3F). On the other hand, at the same stages *msgn1* extended as a ring all around the margin, except on its dorsal aspect, leaving a central gap (Figure 3G-J). For *msgn1* no significant differences were found in the height value at 70% and 80% epiboly (Figure 3K), but similarly to what was observed for *myoD*, the dorsal non-expressing zone was reduced in cavefish embryos at 80% epiboly (Figure 3L). In order to understand the inter-morph differences observed using these two paraxial mesoderm markers, we performed double ISH. Similar to what was observed in single ISH, *msgn1* expression extended further ventrally than *myoD* (Figure 3M, O; compare to insets in Figures 3A, B, G **and** H). Differences also existed in the vegetal to animal axis, where the larger extension encompassed by *myoD* was clear in both morphs (Figure 3M, O). These results suggested that the differences observed in our measurements of paraxial mesoderm extension were mainly due to the cell population expressing *myoD* (but not *msgn1*), which is more advanced towards the animal end of the embryo (Figure 3N, P). In addition, if the size of the central zone where expression of the two paraxial markers is interrupted is taken as readout of dorsal convergence, these data also suggest an earlier convergence and extension in cavefish than in surface fish (at a given stage of epiboly).

**Figure 3.**
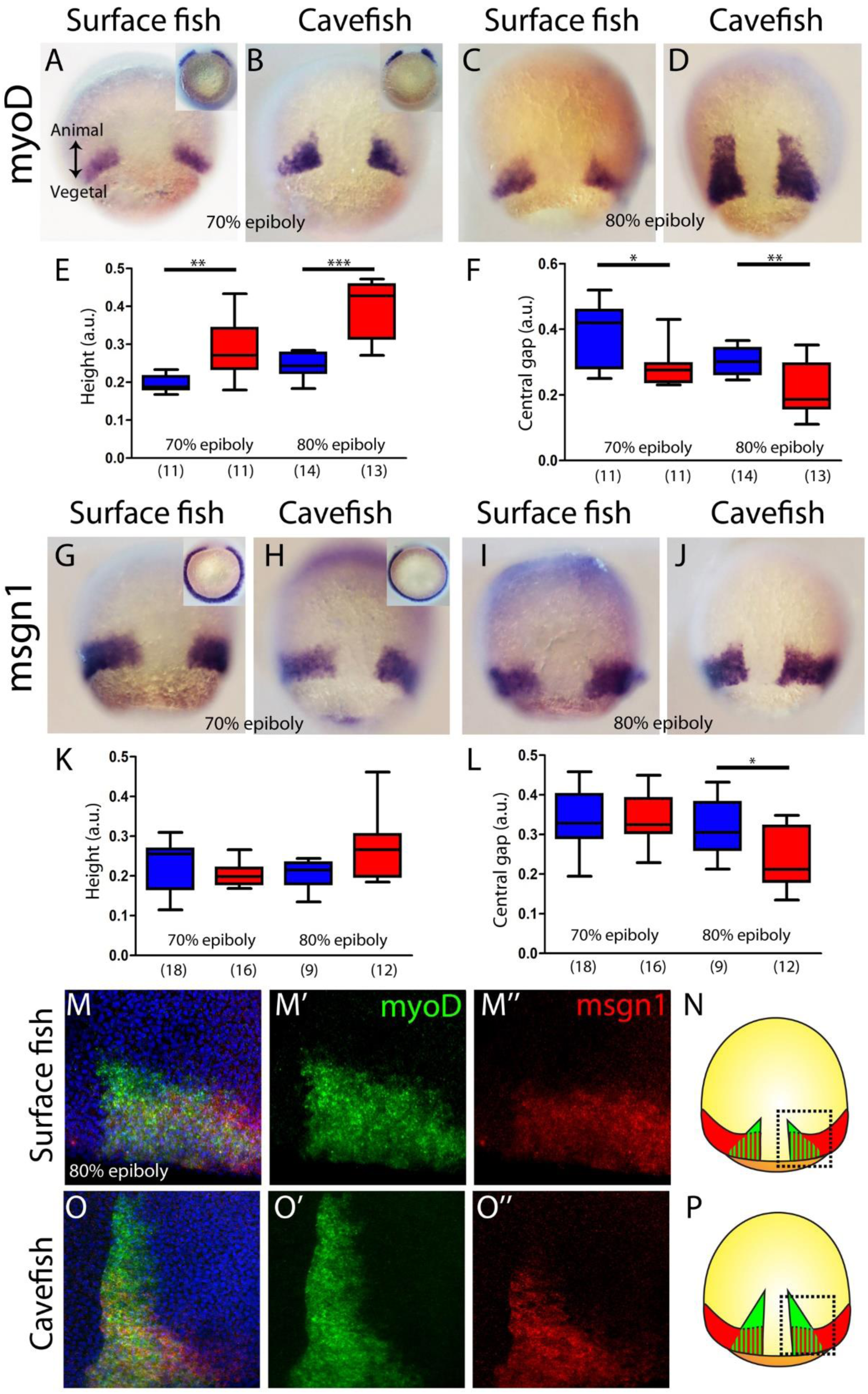
Internalization of paraxial mesoderm in surface fish and cavefish. (A-D) Expression of *myoD* in surface fish (A, C) and cavefish (B, D) at 70 and 80% epiboly (A, B and C, D, respectively). Insets in A and B show the corresponding embryos in a vegetal view. (E) Quantification of height in *myoD* labeled embryos at 70 and 80% epiboly (left and right, respectively). (F) Quantification of central non-expressing zone in *myoD* labeled embryos at 70 and 80% epiboly (left and right, respectively). (G-J) Expression of *msgn1* in surface fish (G, I) and cavefish (H, J) at 70 and 80% epiboly (G, H and I, J, respectively). Insets in G and H show the corresponding embryos in a vegetal view. (K) Quantification of height in *msgn1* labeled embryos at 70 and 80% epiboly (left and right, respectively). (F) Quantification of central non-expressing zone in *msgn1* labeled embryos at 70 and 80% epiboly (left and right, respectively). (M-M’’ and O-O’’). Confocal projection (20-30 µm) showing the expression of *myoD* (green) and *msgn1* (red) double stained surface fish and cavefish embryos (M-M’’ and O-O’’, respectively) at 80% epiboly. DAPI was used as a counterstain (blue nuclei). (N and P) representations of surface fish (N) and cavefish (P) embryos, indicating in black dashed lines the regions of interest showed in M and O. Mann-Whitney tests were performed. ** = 0.0025 (E, left), *** <0.0001 (E, right), * = 0.0181 (F, left), ** = 0.0094 (F, right), * = 0.0209 (L, right). Embryos in dorsal views, animal pole on top; insets on vegetal view, dorsal on top.

### A. mexicanus morphotypes exhibit notable differences in axial mesoderm structure

The antero-posterior embryonic axis in *A. mexicanus* is formed after epiboly has been completed, at the bud-stage (10hpf). The prechordal plate and notochord are the anterior and posterior segments of the axial mesoderm, respectively, both important for the induction and patterning of neural fates. To compare the organization of the axial mesoderm in cave and surface embryos, we analyzed the expression of markers described in the previous sections, to identify specific segments once the antero-posterior axis has been formed. Using triple fluorescent *in situ* hybridization, three non-overlapping molecular subdomains were recognized: the anterior prechordal plate or polster labeled by *dkk1b*, the posterior prechordal plate defined by *shh* expression (wider in cavefish as previously described; Pottin et al., 2011; Yamamoto et al., 2004) and the notochord more posteriorly, labeled by *ntl* (Figure 4A, B). In addition, *lefty1* expression covered both the anterior and posterior subdomains of the prechordal plate (Figure 4C-F). In the posterior prechordal plate *lefty1* and *shh* showed overlapping patterns in both morphotypes (Figure 4C, D), whereas *dkk1b* and *lefty1* showed only minimal co-expression anteriorly (Figure 4E, F), similarly to what we observed at earlier stages (Figure 4 - figure supplement 1). Moreover, the distribution of polster *dkk1b*-expressing cells was strikingly different between the two morphs. In surface fish they were tightly compacted (Figure 4A), while in cavefish they were loosely organized (Figure 4B). The number of *dkk1b*-expressing cells, analyzed in confocal sections, were similar in cavefish and surface fish (Figure 4G). The distribution of the *dkk1b* cells in the antero-posterior axis, measured by the distance between the first and the last cells (Length A-P), was identical (Figure 4H). However, the *dkk1b-*positive cells covered a larger extension in the lateral axis (Length lateral) in cavefish embryos (Figure 4I), indicating that these cells are arranged at a lower density as compared to surface fish. A similar pattern was observed for the anterior domain of *lefty1* expression (compare Figures 4C, E to 4D, F). Thus, both the anterior/polster (*dkk1b*+) and the posterior part (*shh*+) of the prechordal plate are laterally expanded in cavefish.

**Figure 4.**
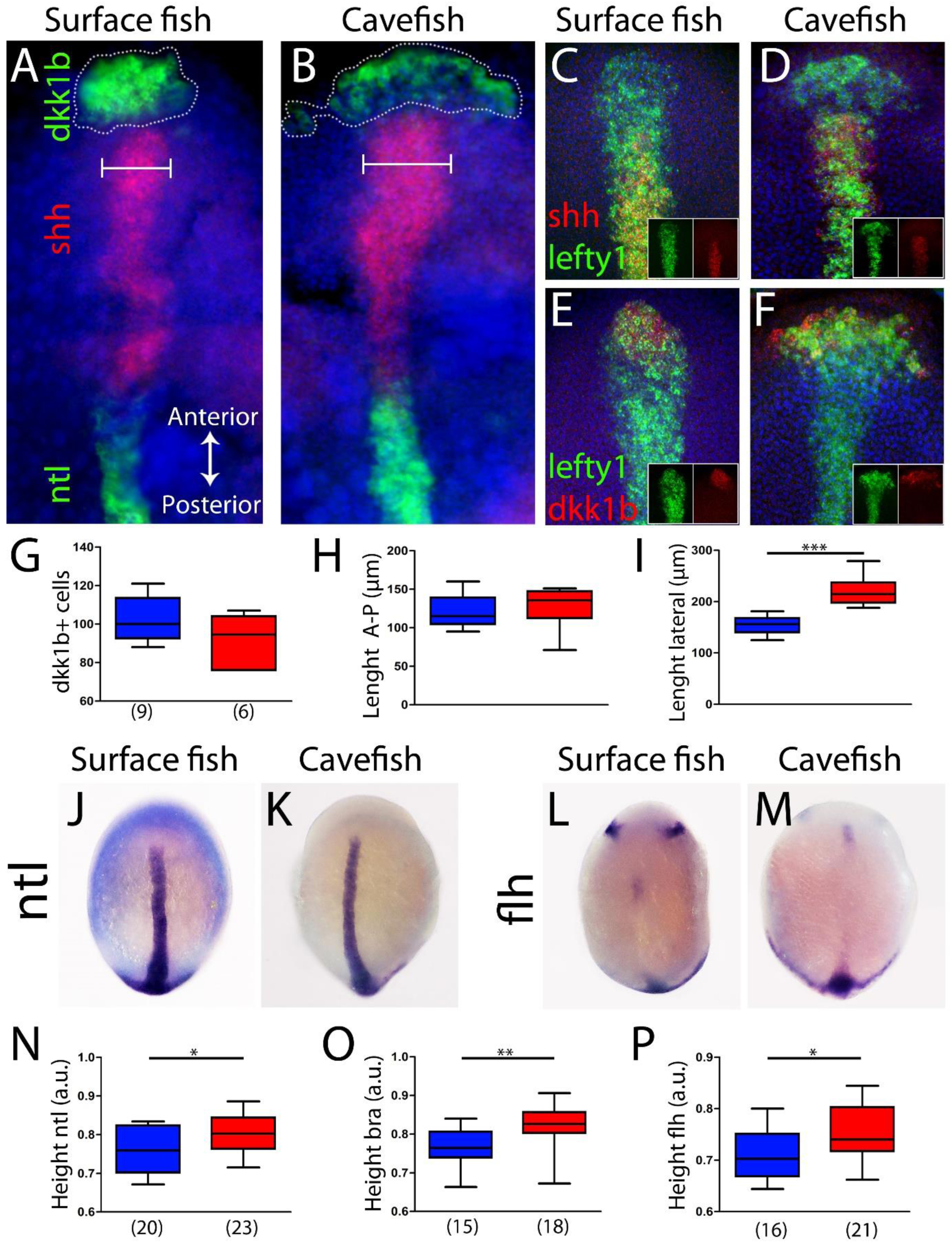
Axial mesoderm organization in surface fish and cavefish. (A-B) Triple ISH to *dkk1b* (green, rostral), *shh* (red, central) and *ntl* (green, posterior) in surface fish (A) and cavefish (B). (C-D) Confocal projection (20-30 µm) showing the expression of *shh* (red) and *lefty1* (green) in surface fish (C) and cavefish embryos (D). Insets show the individual channels. (E-F) Confocal projection (20-30 µm) showing the expression of *dkk1b* (red) and *lefty1* (green) in surface fish (E) and cavefish embryos (F). Insets show the split channels. (G) Quantification of number of cells expressing *dkk1b*. (H) Quantification of the distance between the *dkk1b* expressing cells located in the extremes of the antero-posterior axis. (I) Quantification of the distance between the *dkk1b* expressing cells in lateral extremes. (J-K) Expression of *ntl* in surface fish (J) and cavefish (K). (L-M) Expression of *flh* in surface fish (L) and cavefish (M). (N) Quantification of height in *ntl* labeled embryos. (O) Quantification of height in *bra* labeled embryos. (P) Quantification of height in *flh* labeled embryos. All embryos at tail bud stage, in dorsal view, anterior upwards. Pictures A-F are flat mounted embryos and pictures J-M are whole mount embryos. Mann-Whitney tests were performed. *** < 0.0001 (I), * = 0.0396 (N), ** = 0.0012 (O), * = 0.0142 (P).

Next, other differences in size or position of axial mesoderm segments at bud stage were explored. The distance from the anterior-most polster cell expressing *dkk1b* to the leading notochord cell expressing *ntl* was identical in the two morphs (Figure 4 - figure supplement 2A-C). Polster cells expressing *dkk1b* laid just beneath the cells of the anterior neural plate border (*dlx3b*+) in both morphotypes (Figure 4 - figure supplement 2D, E). The extension of the notochord was also measured. At bud stage *ntl* and *bra* expression labeled the notochord in its whole extension (Figure 4J, K and not shown). On the other hand, *flh* was expressed in the posterior end and in a small cluster of the rostral notochord (Figure 4L, M) (plus two bilateral patches in the neural plate probably corresponding to the prospective pineal gland in the diencephalon). For the three notochordal markers, the distance from the rostral expression boundary to the tail bud (normalized by the size of the embryo) was larger in cavefish compared to surface fish (Figure 4N-P). In line with our observations of axial and paraxial mesoderm markers during mid-gastrulation (Figure 2-3), these results suggest a precocious convergence and extension in cavefish compared to surface fish.

### Testing the effects of heterochrony in gastrulation and gene expression dynamics on brain development

In zebrafish embryos *dkk1b* expression in the prechordal plate becomes downregulated from early somitogenesis (Hashimoto et al., 2000). Our observations of heterochronic gastrulation events prompted us to search for potential differences also in the disappearance of *dkk1b* expression later on. In surface fish *dkk1b* was still expressed in all embryos at the 6 and 8 somite stage (13/13, not shown and 17/17, Figure 5A, 6E **right**). In contrast, in cavefish *dkk1b* expression was observed only in 46% of the embryos at 6 somites (6/13, always with low signal level) (not shown) and it was absent in 64% of embryos at 8 somite stage (21/33, Figure 5B **and** 6E **right**).

**Figure 5.**
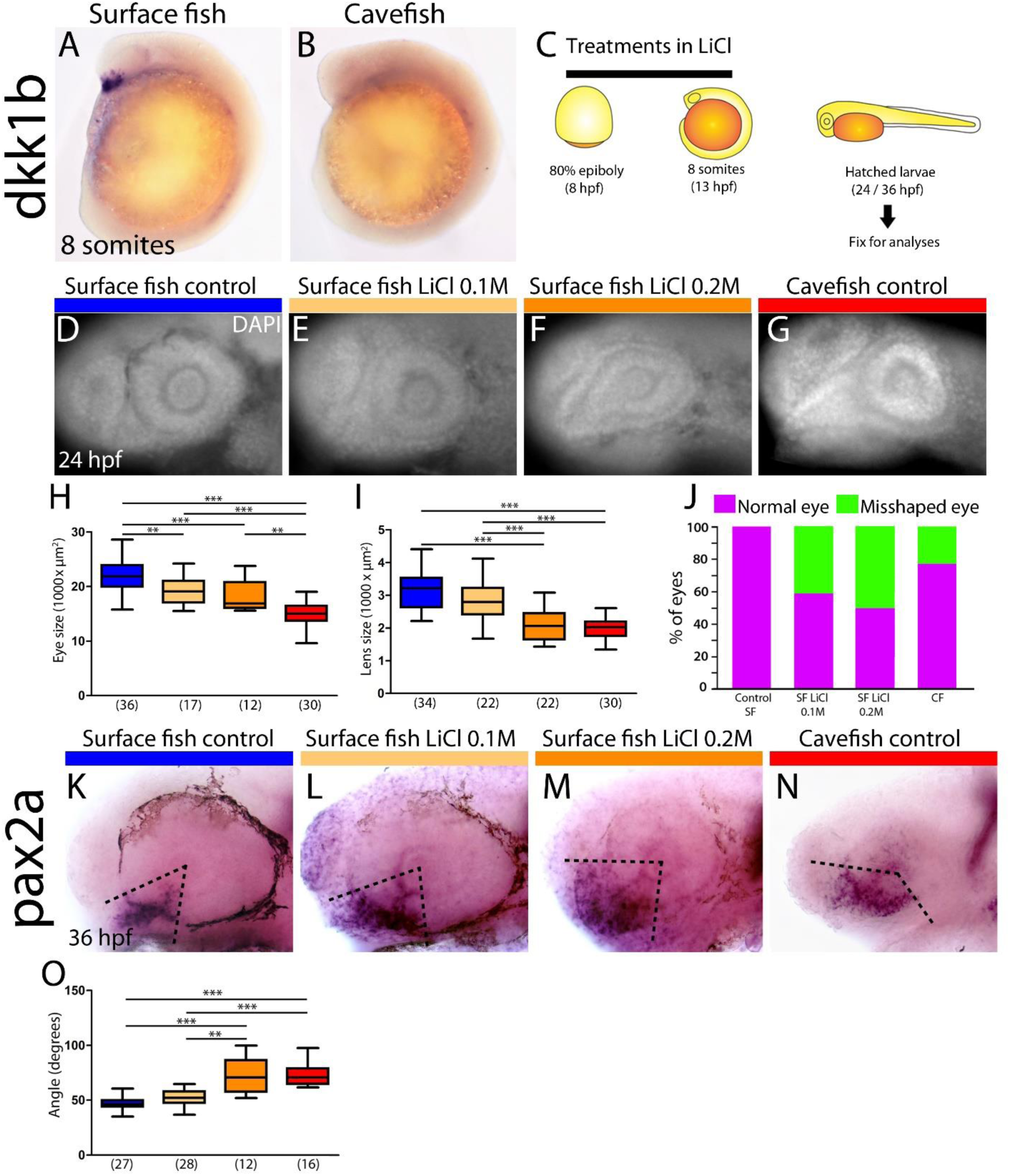
Differential off-set of *dkk1b* expression may be relevant for the optic phenotype in cavefish. (A-B) Expression of *dkk1b* at 8 somite stage in surface fish (A) and cavefish (B). (C) Experimental design for LiCl treatments. Dechorionated surface fish embryos were treated in LiCl solutions from the end of gastrulation (8hpf, left) until mid-somitogenesis (13hpf, center) and then fixed for analyses at larvae stages (24 or 36hpf, right). (D-J) Effect of LiCl treatments analyzed at 24hpf. (D-E) Surface fish untreated (D), treated with LiCl 0.1 and 0.2M (E and F, respectively) and cavefish untreated (G), stained with DAPI at 24hpf. Quantification of the eye size (H) lens size (I) and percentage of embryos with misshaped developing eye (J). (K-O) Effect of LiCl treatments analyzed at 36hpf. Expression of *pax2a* at 36 hpf in the optic stalk/optic fissure in surface fish untreated (K), treated with LiCl 0.1 and 0.2M (L and M, respectively) and cavefish untreated (N). Quantification of the angle measured indicated in K-N in black dashed lines. Kruskal-Wallis tests with Dunn’s post-test, were performed.

**Figure 6.**
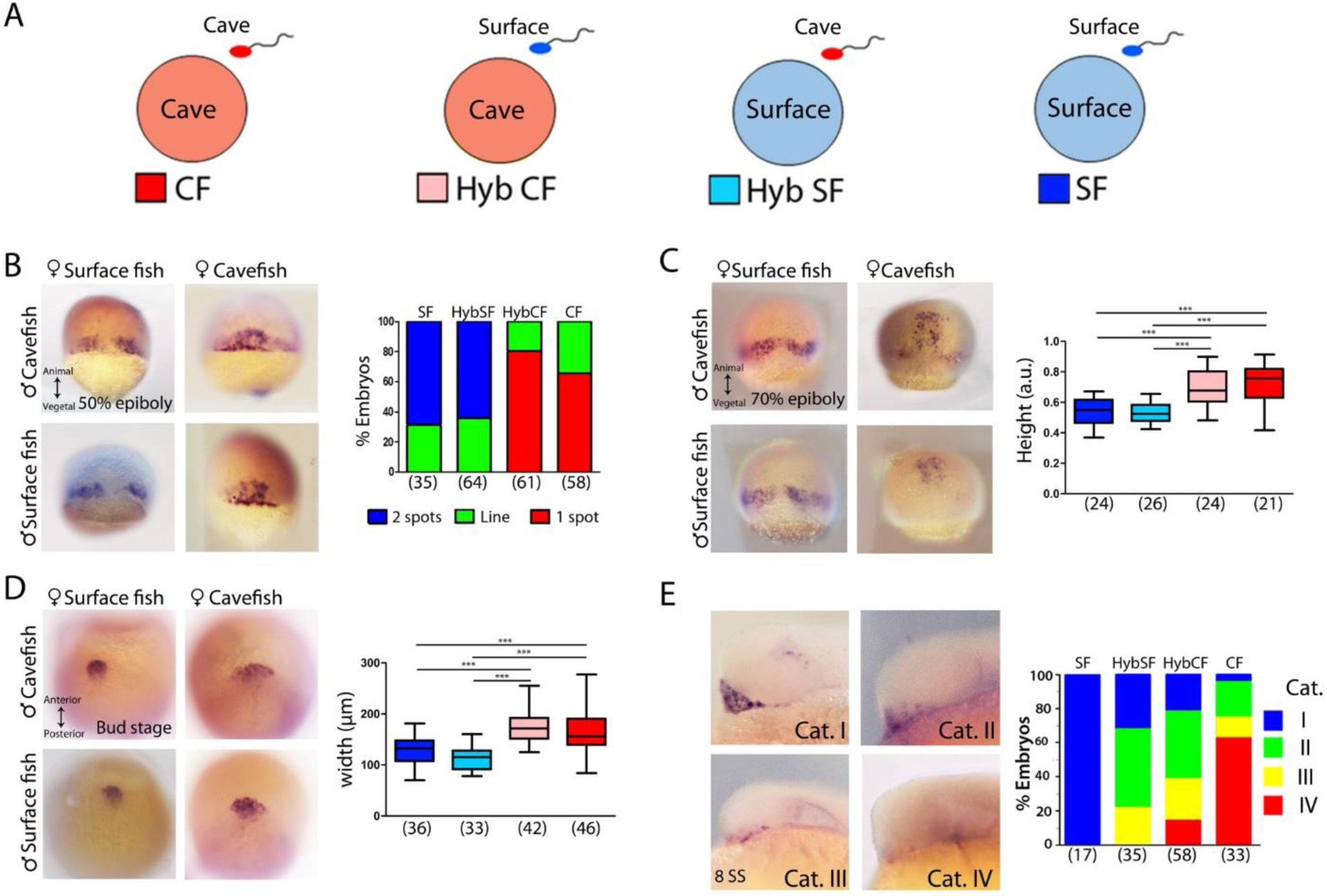
Maternal effect on early development. Schematic representation of the fertilizations performed for the analyses of maternal effect in F1 hybrids. Oocytes from either morph (cave in pink and surface in light blue) were fertilized with sperm coming from cavefish (red) or surface fish (blue). For simplicity, F1 hybrids were named HybCF (oocyte from cavefish, pink) and HybSF (oocyte from surface fish, light blue), based on their maternal contribution. (B-E) Expression of *dkk1b* at 50% of epiboly (B), 70% of epiboly (C), bud stage (D) and 8 somite stage (E). In the panels B-D are shown HybSF (top left), cavefish (top right), surface fish (bottom left) and HybCF (bottom right). Quantification of the expression pattern of *dkk1b* at 50% epiboly (B, right), classified into three categories: “Two spots” (blue) is the pattern observed in the panel on the left column, “1 spot” (red) is the pattern observed in the panel on the right column, and “Line” is an intermediate profile (not shown). The Y-axis indicates the percentage of the total embryos belonging to each of the categories and the numbers of embryos examined are indicated. Quantification of height in *dkk1b* labeled embryos at 70% epiboly (C, right). Quantification of width of the polster based on *dkk1b* expression (D, right). (E, right) Quantification of the pattern of *dkk1b* at 8 somite stage, where embryos were classified according to the number of positive cells. Category I (blue, in the panel embryo on top left, surface fish), more than 5cells; category II (green, in the panel embryo on top right, HybSF), between 3 and 5 cells; category III (yellow, on the panel bottom left, HybCF); and category IV, no positive cells (red, on the panel bottom right, cavefish). All embryos imaged in whole mount, embryos in B and C dorsal view with animal pole upwards, embryos in D in dorsal view with anterior upwards and embryos in E in lateral view with anterior to the left. Kruskal-Wallis tests with Dunn’s post-test, were performed in all cases.

Given the major spatio-temporal differences in *dkk1b* expression pattern observed from the onset of gastrulation to the end of neurulation between cave and surface embryos, we also examined its expression levels through qPCR. While at 50% epiboly *dkk1b* transcript levels were similar in the two morphs (0.95 fold, NS), at bud stage *dkk1b* levels were almost four times lower in cavefish than in surface fish embryos (0.27 fold).

Since *dkk1b* is a strong inhibitor of Wnt signaling, with conserved functions in the regulation of brain development (Hashimoto et al., 2000; Lewis et al., 2008), the observed differences in cellular arrangement, expression levels and timing of downregulation in the two *Astyanax* morphotypes may have downstream consequences in forebrain morphogenesis. This hypothesis was partly tested by treating surface fish embryos with LiCl (Figure 5C), a Wnt-βcat pathway activator, to mimic the cavefish situation in which the Wnt antagonist *dkk1b* is expressed at lower levels. In line with results reported in zebrafish (Shinya et al., 2000), LiCl treatments (0.1 and 0.2M, from 8 to 13hpf) produced a decrease of the size of the optic vesicle in SF at 13hpf (not shown), and a reduction of the size of the retina and lens at 24hpf (Figure 5D-F, H, I), which are hallmarks of cavefish embryonic eye morphology (Yamamoto et al 2004; compare to Figure 5G). In addition, manipulation of the levels of Wnt-βcat signaling in surface fish produced a misshaped retina with a wider optic stalk (Figure 5F). This was observed in 41% and 50% of the examined eyes of embryos treated with LiCl 0.1M and 0.2M, respectively (Figure 5J). A similar phenotype was seen in 23% of cavefish embryos at the same stage (Figure 5G, J). The interpretation of this morphological phenotype was confirmed molecularly, as the expression domain of the optic stalk marker *pax2a* was significantly wider at 36hpf in surface fish embryos exposed to LiCL and in cavefish embryos than in untreated animals (Figure 5K-N, O).

Together these data strongly suggest that modified levels of Wnt signaling during early embryogenesis might participate to the developmental evolution of cavefish eye defects.

### Maternal determinants influence early developmental differences in A. mexicanus morphotypes

The earliest developmental events, including the first cell divisions, breaking of symmetries and induction of the embryonic organizer, rely exclusively on maternal factors deposited in the oocyte before fertilization. The findings above showing earlier convergence, extension and internalization of mesodermal cell populations in the cave morphs, together with differences in the spatio-temporal gene regulation in tissues derived from the organizer, prompted the examination of precocious embryogenesis and the investigation of maternal components. The inter-fertility between *A. mexicanus* morphotypes offers a powerful system to study the potential contribution of these maternally-produced factors to phenotypic evolution (Ma et al., 2018). We compared gastrulation progression in F1 hybrid embryos obtained from fertilization of surface fish eggs with cavefish sperm (HybSF), and cavefish eggs with surface fish sperm (HybCF) (Figure 6A). In principle, phenotypic correspondence to the maternal morphotype indicates a strong maternal effect. Results obtained in F1 hybrids were compared to those obtained from wild type morphs in previous sections.

First, the expression patterns of *dkk1b* during development were compared. At 50% epiboly the percentages of the phenotypic categories (described in Figure 1A-C) in hybrid embryos were strikingly similar to those of their maternal morphotypes, with the majority of HybSF presenting two spots of *dkk1b* expression as surface fish embryos, whereas most of HybCF embryos showed only one continuous expression domain (Figure 6B). At 70% epiboly the results followed the same trend. In HybSF embryos the two domains of *dkk1b* expressing cells start to join dorsally, with little advancement towards the animal pole, similar to surface embryos (Figure 6C). In contrast, HybCF were more alike cavefish embryos, with cells grouped dorsally close to the animal end (Figure 6C). Analyses of the distance reached by the leading cell showed significant differences between the two reciprocal hybrids types, which were identical to their maternal morphs (Figure 6C**, right**). The expression of *lefty1* and *ntl* at 70% epiboly was also examined in F1 hybrids (Figure 6 - figure supplement A and B, respectively). The advancement of axial mesoderm populations labeled by the two markers was significantly increased in HybCF compared to HybSF, with height values akin to their respective maternal morphs (Figure 6 - figure supplement A, **right and** B, **right**). These results indicate that spatio-temporal differences observed during gastrulation between cavefish and surface fish fully depend on maternal contribution.

In *A. mexicanus* the prechordal plate at the end of gastrulation showed marked morphotype-specific differences in cell organization. We evaluated the impact of maternal determinants on these differences by comparing the expression of *dkk1b* in the F1 hybrids with that of their parental morphotypes (Figure 6D). We found a broader distribution of *dkk1b-*expressing cells in the HybCF (Figure 6D**, center bottom**) compared to HybSF (Figure 6D**, left bottom**). The patterns observed in the F1 hybrids were identical to the patterns on their maternal morphs (Figure 6D**, right**), highlighting the effect of the oocyte composition up to the end of gastrulation.

Next, we tested the maternal contribution to the disappearance of *dkk1b* expression during mid-somitogenesis described in the previous section. At the 8 somite stage, the segregation of phenotypes in reciprocal F1 hybrids was not as clear as in the parental morphs (Figure 6E). For this reason, we decided classify the expression patterns of *dkk1b* in four distinct categories: I, widely expressed in the prechordal plate (Figure 6E**, top left; blue**); II, clear expression in 3-5 cells (Figure 6E**, top right; green**); III, clear expression in 1-2cells (Figure 6E**, bottom left; yellow**); and IV, absence of expression (Figure 6E**, bottom right; red**). In hybrids, we found similar percentages of intermediate categories II and III (63-64% in both cases). However, in HybCF there was an important proportion of category IV embryos (no expression, 16%), closer to the cavefish, while none of the HybSF fell in this category, like surface fish embryos. From this result we deduced that the downregulation of *dkk1b* expression is still under the influence of maternal factors, although this influence is weaker than at earlier stages.

Finally, we sought to test whether forebrain phenotypes previously described in cavefish at later embryonic stages could be also influenced by maternal effects, as a long-lasting consequence of the maternal influence during gastrulation. Hypothalamic, eye and olfactory epithelium development were analyzed in reciprocal F1 hybrids between 15hpf and 24hpf.

Inter-morph variations in the expression domains of the LIM-homeodomain transcription factors *Lhx9* and *Lhx7* drive changes in Hypocretin and NPY neuropeptidergic neuronal patterning in the hypothalamus, respectively (Alié et al., 2018). We therefore compared expression domains of *Lhx9* (size of the hypothalamic domain at 15 hpf; brackets in Figure 7A**, left**) and *Lhx7* (number of positive cells at 24 hpf in the hypothalamic acroterminal domain; dotted circles in Figure 7B**, left**) and the numbers of their respective neuropeptidergic Hypocretin and NPY derivatives in the reciprocal hybrids and their parental morphotypes (Figures 7C **and** D, respectively). In all four cases the analyses showed strong significant differences between cavefish and surface fish, as previously described (Alié et al., 2018) (Figure 7A-D histograms, *** for each). In order to help the visualization and interpretation of the F1 hybrid data, simplified plots were generated (Figure 7A-D**, right**) with the mean values for cavefish and surface fish in the extremes (red and blue dots, respectively), an average black dot representing the expected value for the phenotype if there is no effect of any kind (maternal, paternal or allelic dominance), and the HybSF and HybCF values (light blue and pink respectively). If experimental values are closer to the maternal morphotype, it can be interpreted as the phenotype being under maternal regulation. Other possibilities, such as a mix of maternal and zygotic influence, or recessive or dominant effects in heterozygotes can also be interpreted.

**Figure 7.**
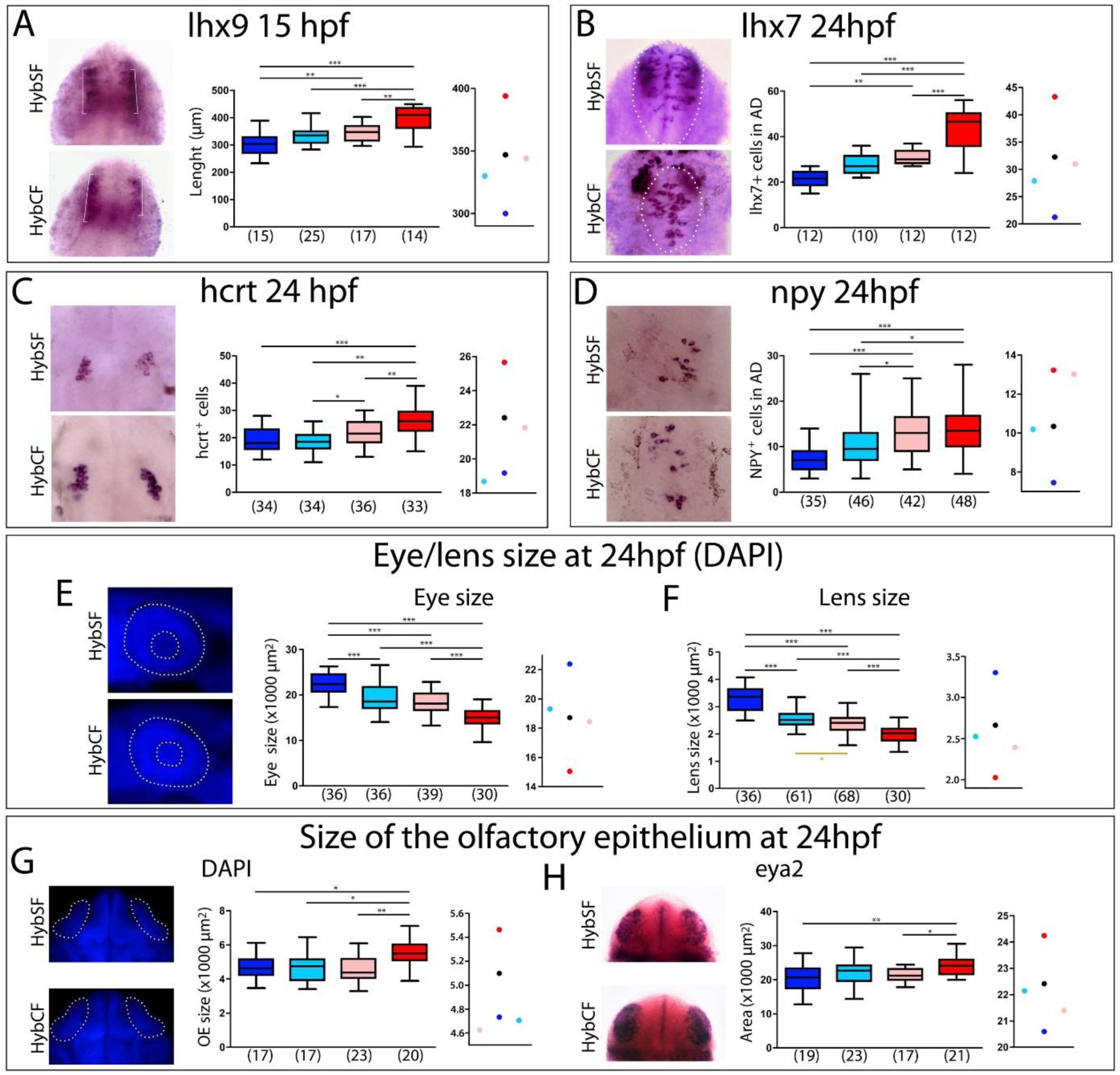
Maternal effect on later embryogenesis. (A) Expression of *lhx9* in HybSF (left, on top) and HybCF (left, on bottom) at 15 hpf. Quantification of the length of the expression domain in the prospective hypothalamus (white brackets) (center) and the corresponding plot of means distribution (right). (B) Expression of *lhx7* in HybSF (left, on top) and HybCF (left, on bottom) at 24 hpf, with the acroterminal domain indicated in dashed lines. Quantification of the number of *lhx7* expressing cells in the acroterminal domain (center) and the corresponding plot of means distribution (right). (C) Expression of *hcrt* HybSF (left, on top) and HybCF (left, on bottom) at 24 hpf. Quantification of the number of hypothalamic *hcrt* expressing cells (center) and the corresponding plot of means distribution (right). (D) Expression of *NPY* in HybSF (left, on top) and HybCF (left, on bottom) at 24 hpf in the acroterminal domain. Quantification of the number of *NPY* expressing cells (center) and the corresponding plot of means distribution (right). (E) DAPI stained (left, on top) and HybCF (left, on bottom) embryos at 24 hpf. White dashed lines indicate the contour of the eye ball and the lens (exterior and interior circles, respectively). Quantification of the size of the Eye ball (center) and the plot of means distribution (right). (F) Quantification of the size of the lens (left), and plots of means distribution (right). (G) DAPI stained HybSF (left, on top) and HybCF (left, on bottom) embryos at 24 hpf. White dashed lines indicate the contour of the olfactory placodes. Quantification of the size of olfactory placodes (center) and the corresponding plot of means distribution (right). (H) Expression of *eya2* in HybSF (left, on top) and HybCF (left, on bottom) at 24 hpf. Quantification of the size of the *eya2* expression domain in the olfactory placodes (center) and the corresponding plot of means distribution (right). Embryos in A-D were dissected and mounted in ventral view, anterior upwards. Embryos in E-H are whole mounted embryos in lateral view anterior to the left (E); in dorsal view, anterior upwards (G); or in frontal view, dorsal upwards (H). Kruskal-Wallis tests with Dunn’s post-test, were performed in all cases.

For *Lhx9* and *Lhx7,* hybrids values were similar and intermediate between the cave and surface morphs, with a slight deviation towards the surface morph, more evident for the HybSF (Figure 7A **and** B, **center and right**). For Hypocretin and NPY neuropeptidergic lineages derived from *Lhx9* and *Lhx7-*expressing progenitors, respectively, a significant difference in neuron numbers existed between reciprocal hybrids (Figure 7C **and** D, **\*** for each), suggesting the involvement of maternal components. Moreover, the number of Hypocretin neurons in HybSF and the number of NPY neurons in HybCF were identical to their maternal morphotype, whereas values for their reciprocal hybrids were close to the theoretical intermediate value (Figure 7C **and** D, **on the right**). These results suggest that maternal determinants impact at least in part hypothalamic neuronal differentiation, possibly together with other, complex, allelic dominance or zygotic mechanisms.

In cavefish, the smaller size of the eye primordium and the larger olfactory epithelia compared to surface fish are also due to modifications of signals emanating from midline organizers, including Shh and Fgf8 (Hinaux et al., 2016; Pottin et al., 2011; Yamamoto et al., 2004). The size of these sensory structures were measured in reciprocal hybrids to test if the cascade of events affected by the maternal determinants also has an impact in their later development. First, in F1 hybrids, the size of the eye ball and the size of the lens at 24 hpf (dotted lines in Figure 7E, DAPI stained embryos) were intermediate between those from the parental morphs (Figure 7E **and** 7F**, respectively**), without significant differences between the hybrids in the ANOVA test. Of note, when considering only the hybrids, the Mann Whitney test showed a significant difference in lens size (Figure 7F, golden star p= 0.0202), suggesting a reminiscence of maternal effect. In the plot of mean distribution for eye ball size the F1 hybrid values were close to the expected mean (Figure 7E**, right**). In the plot for lens size however, the hybrid values were slightly deviated towards the cavefish mean (Figure 7F**, right**), suggesting also a dominance of cavefish alleles involved in lens development. Finally, the size of the olfactory epithelium at 24 hpf was similar in HybSF and HybCF (Figure 7G; DAPI staining, and Figure 7H; ISH to *eya2*). In both types of read-outs, the mean values for hybrids appeared shifted towards surface fish values, suggesting a dominance of the surface fish alleles involved in the development of the olfactory epithelium.

Taken together, these results indicate that the effect of maternal determinants are fully penetrant up top final stages of gastrulation, suggesting that RNAs and proteins present in the oocyte must vary between the two *Astyanax* morphotypes. At later developmental stages the maternal effect appears to be “diluted” by other mechanisms regulating gene expression and morphogenesis, although some differences can still be observed.

### Towards identification of varying maternal factors in cavefish

To obtain an exhaustive molecular view of maternal transcriptomic differences between surface and cavefish, RNA-sequencing was performed on *Astyanax* embryos at 2-cell stage (surface fish, n=2 samples; cavefish, n=3 samples; and reciprocal F1 hybrids, n=3 samples each). The dataset (between 75 and 100 million paired reads per sample) was analyzed through the European Galaxy Server and reads were aligned to the Surface Fish *Astyanax* genome (NCBI, GCA_000372685.2 Astyanax_mexicanus-2.0). The sample-to-sample distance analysis grouped the four types of samples in two clear categories, strictly depending on their maternal contribution (Figure 8A). Similarly, Principal Component Analyses (PCA) analyses clustered the samples from hybrid embryos together with those coming from their maternal morphotype (Figure 8 - figure supplement A). These results clearly confirmed that the paternal contribution has no influence on the egg transcriptome at this stage, so we decided to combine the samples according to their mother morphotype (pooled surface fish and hybSF, *versus* pooled cavefish and hybCF), thus increasing the number of samples per condition, and rendering downstream analysis easier and more powerful. To quantitatively compare the transcriptome of cave and surface eggs, the numbers of differentially expressed genes (DEG) were assessed (see Methods). Among the 20730 genes that were expressed at 2-cell stage, close to a third (32%) were differentially expressed between surface and cavefish (Figure 8B). A similar proportion was up- or down-regulated in cavefish relative to surface fish (17.25% and 14.69%, respectively). To get insights on which biological functions were the most different between eggs of the two morphotypes, a gene ontology (GO) enrichment analysis was carried out on DEGs with an absolute fold change higher than 5 (log(FC)> 2.32193). Among the significantly enriched biological processes that might be most relevant for our work we found cell adhesion (7.1%) and signaling (6.5%) (Figure 8C). When analyzing separately up- and down-regulated genes for GO enrichment, no biological process was found enriched in down-regulated genes, whereas the above mentioned processes were still found enriched in the up-regulated gene subset (Figure 8 - figure supplement B). This means that genes involved in ion transport, cell adhesion and cell signaling are mainly up-regulated in cavefish eggs compared to surface fish eggs. It is also worth noting that genes involved in metabolism show significant enrichment when analyzing all the DEGs (fold change higher than 1.5), meaning that “metabolic” transcripts mostly show fold changes lower than 5 (not shown). Hence the most strongly dysregulated genes are not the ones involved in metabolism but those involved in signaling and cell interactions. Together, these results show that the RNA composition of the cavefish and surface fish eggs shows a strong maternal signature, and thus oocyte content could contribute to the developmental evolution of cavefish phenotype.

**Figure 8.**
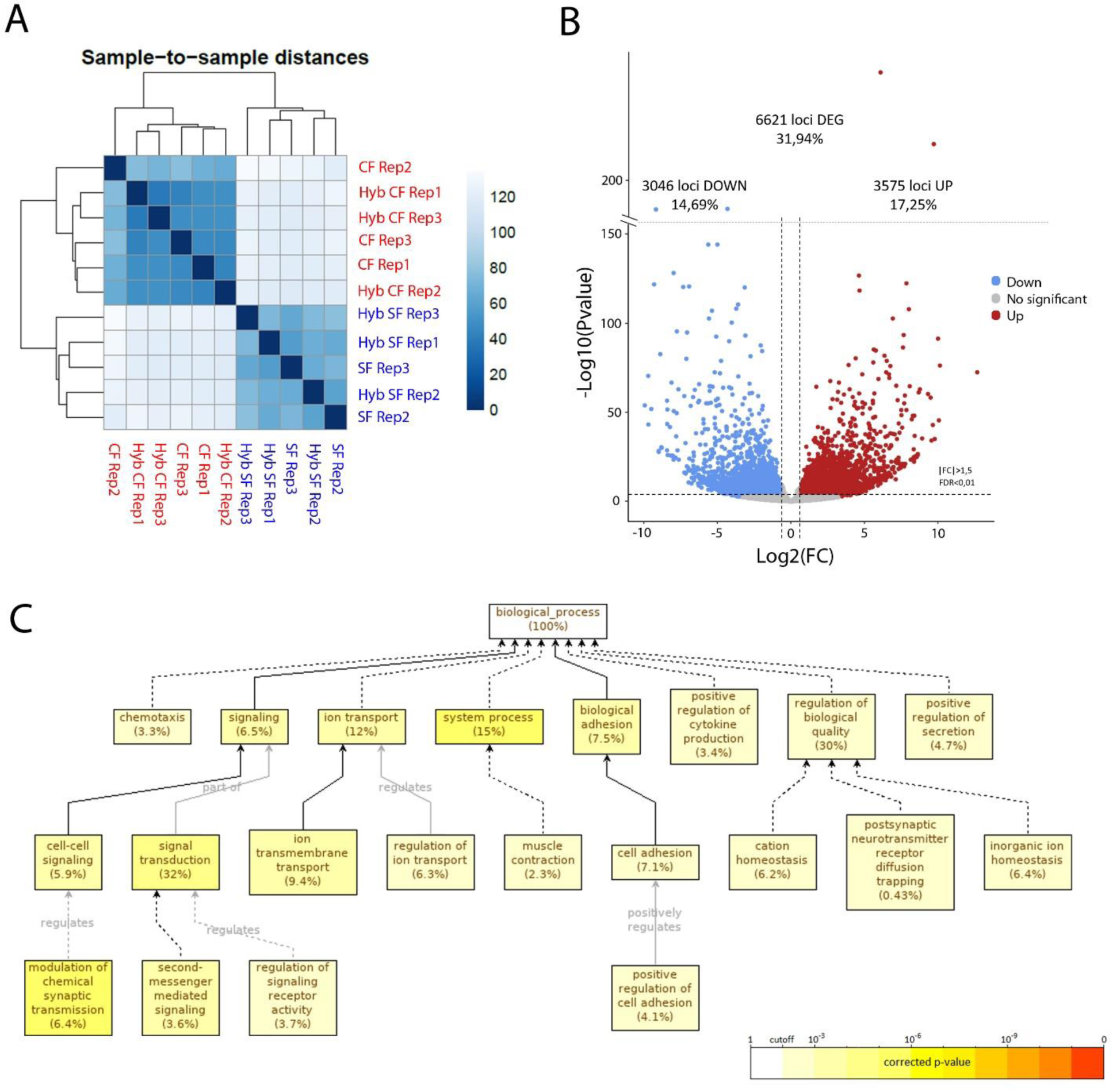
RNA-sequencing of maternal mRNA of surface fish (SF), cavefish (CF) and reciprocal F1 hybrid (HybSF and HybCF, respectively) eggs at 2-cell stage. (A) Sample-to-sample distance between all samples. Samples that are similar are close to each other. In the scale, lower numbers (dark blue) indicate closer relationship between samples than higher numbers (light blue/white). (B) Volcano plot of expressed genes at 2-cell stage (n=20730). Genes with an absolute fold change >1.5 and an adjusted p-value (FDR) <0.01 are considered differentially-expressed in cavefish compared to surface fish. Up-regulated genes in cavefish are in red and down-regulated genes in cavefish genes are in blue. (C) Gene ontology enrichment (level: Biological Process) for cavefish DEGs with an absolute fold change higher than 5. Black lines correspond to “is a” relationship, whereas grey lines correspond to the annotated relationship. Full lines correspond to direct relationship and dashed lines to indirect relationship (i.e. some nodes are hidden). The color of a node refers to the adjusted p-value (FDR) of the enriched GO term and the percentage corresponds to the frequency of the GO term in the studied gene set at the level considered. A given gene can have several GO terms. Only enriched GO terms that pass the threshold (p-value<0.01) are displayed on the graph.

Finally, we picked two candidate genes from the transcriptomics dataset, that were directly relevant to our findings in the previous section: *Oep* (*one-eyed pinhead,* also named *tdgf1*), a Nodal co-receptor necessary for *dkk1b* induction and shield formation and whose maternal and zygotic mutant (*MZoep*) shows defects in margin internalization and fate specification in zebrafish (Carmany-Rampey & Schier, 2001; Zhang et al., 1998); as well as the maternal ventralizing transcription factor *Vsx1* (*Visual System homeobox 1*) which regulates *flh* and *ntl* expression and is involved in axial *versus* paraxial mesoderm specification and migration (He et al., 2014; Xu, He et al, 2014). qPCR analyses, on 2hpf embryos, showed that *Vsx1* and *Oep* mRNA levels were significantly reduced in cavefish (2.50 and 1.75 times less expressed in cavefish, respectively) confirming the RNA-seq results (8.21 and 1.63 times less respectively). To test for a possible role of these two maternal down-regulated transcripts in the cavefish gastrulation phenotype, we performed overexpression experiments through mRNA injection at one cell stage in cavefish eggs. As read-out of these rescue experiments, *dkk1b* expression was examined at 50% and 70% epiboly. *Vsx1*-injected and *Oep*-injected embryos were similar to control cavefish embryos in terms of spatio-temporal *dkk1b* pattern, although some signs of disorganization were visible on several specimens (not shown). Thus, a role for *Vsx1* and *Oep* maternal transcripts in the variations of *dkk1b* expression observed between the two *Astyanax* morphs is unlikely. Future experiments should focus on transcripts showing high fold-changes of expression between cavefish and surface fish.

## Discussion

*Astyanax mexicanus* has become an excellent model to uncover developmental mechanisms leading to phenotypic evolution. Modifications in midline signaling centers during early embryogenesis have led to troglomorphic adaptations in cavefish, including eye degeneration, larger olfactory epithelia and increased number of taste buds. Here we show striking temporal, spatial and quantitative differences in the expression of the Wnt inhibitor *dkk1b* at shield stage and during gastrulation, and we explore the idea that maternally-regulated gastrulation might be a source of variation contributing to cavefish morphological evolution.

### Prechordal plate and forebrain patterning

Genetic manipulations, tissue ablation and transplantation experiments have demonstrated the importance of the prechordal plate as a signaling center involved in the patterning of the basal forebrain (Heisenberg & Nüsslein-Volhard, 1997; Pera & Kessel, 1997). In fish, the prechordal plate is organized in two domains: the rostral polster (Kimmel et al., 1995) and a posterior domain, abutting caudally with the notochord. In *A. mexicanus* the expression of *shh* in the posterior prechordal plate occupies a wider domain in the cavefish compared to surface fish (Pottin et al., 2011; Yamamoto 2004), and enhanced shh signaling has pleiotropic effects in in the development of head structures in the cavefish (Yamamoto et al., 2009). Here we showed that the anterior domain of the prechordal plate is a source of the morphogen dkk1b, whose expression is complementary to that of *shh* at the neural plate stage (Figure 4A-B). At this stage, *dkk1b* expressing cells are organized as a compact cluster in surface fish, while in cavefish they are more loosely distributed, and with lower levels of *dkk1b* transcripts. Inhibition of Wnt signaling in the presumptive anterior brain is critical for patterning and morphogenesis. Mouse or *Xenopus* embryos with impaired Dkk1 function lack anterior brain structures (Glinka et al., 1998; Mukhopadhyay et al., 2001), whereas misexpression of *dkk1b* in zebrafish embryos produce anteriorization of the neurectoderm, including enlargement of eyes (Shinya et al., 2000). In *Astyanax* also, we found that Wnt activation in surface fish embryos by LiCl-treatments, phenocopying the naturally occurring cavefish condition where lower levels of *Dkk1b* transcripts could lead to lower Wnt inhibition, leads to a reduction of eye and lens size (Figure 7C-J) and an expansion of optic stalk tissue (Figure 7K-O), both cavefish-specific hallmarks of eye development (Yamamoto et al., 2004; Devos et al., 2018). Head development is sensitive to Wnt signaling dosage (Lewis et al., 2008), and the temporal variations of *dkk1b* expression we observed here might contribute to forebrain evolution in cavefish. Indeed, the timing and intensity of Wnt (this work) and Bmp (Hinaux et al., 2016) signaling at the anterior pole of the axial mesoderm must instruct the fate and morphogenetic movements of overlying anterior neural plate progenitors destined to form the optic region and the hypothalamus, as well as the placode derivatives (Bielen et al., 2017; Rétaux et al., 2013).

### Embryonic axis formation

The establishment of the embryonic axes and primordial germ layers occurs through complex morphogenetic cell rearrangements during gastrulation (Schier and Talbot, 2005; Solnica-Krezel and Sepich, 2012). The main outcomes of gastrulation are the spreading of the blastodermal cells, internalization of endomesoderm precursors and the elongation of the antero-posterior embryonic axis. We hypothesized that the differences observed in the axial mesoderm of *A. mexicanus* morphotypes may be the consequence of upstream events during gastrulation. At equivalent stages, as judged by the percentage of epiboly, we observed that the advancement of internalized tissues migrating in the vegetal to animal direction is more precocious in cavefish embryos than in surface fish. Interestingly, this finding was not only restricted to axial mesodermal elements, but also applied to laterally adjacent paraxial mesoderm, suggesting a global phenomenon. From the different measurements performed, we inferred that dorsal convergence and anteroposterior extension might be the driving forces leading to the more advanced phenotype observed in cavefish gastrulas. Interestingly, the differences in hypoblast movements we observed, relative to the percentage of epiboly, highlight the uncoupling between gastrulation cell movements and the epiboly itself, as spectacularly illustrated in the extreme example of annual killifish embryogenesis (Pereiro et al., 2017). We suggest that these temporal variations in gastrulation events might later correlate to differences observed in the off-set of *dkk1b* expression, starting in cavefish before the 6 somite stage and in surface fish after the 8 somite stage.

### Cellular interactions during gastrulation

Gastrulation involves dynamic interactions between different cell populations, while as they move, cells are exposed to changing signals in their immediate environment. Individual interactions between tissues, such as the migration of the hypoblast using epiblast as substrate (Smutny et al., 2017) and the influences that the blastodermal cells receive from direct physical contact with the extraembryonic EVL (Reig et al., 2017) and YSL (Carvalho & Heisenberg, 2010) must be integrated as gastrulation proceeds. In addition, the prechordal plate has been described as a cell population undergoing collective migration, implying numerous cell-cell interactions between prechordal cells themselves (Dumortier et al., 2012; T. Zhang et al., 2014). Genetic dissection of the parameters regulating prechordal plate migration (Kai et al., 2008), as well as the identification of intrinsic properties of the moving group (Dumortier et al., 2012), have helped understanding the molecular and cellular aspects regulating their migration. The markers we used here to label the prechordal plate during gastrulation suggest that within this domain *lefty1*-expressing cells follow collective migration as a cohesive group, whereas *dkk1b*+ cells constitute a more dispersed group, especially in the cavefish, and as also recently observed by Ren et al., 2018. Moreover, increased Nodal signaling and changed cell distribution have been reported in the organizer in cavefish embryos (Ren et al., 2018). Together with our observation of earlier movements of axial mesoderm cells in cavefish, these data suggest that the structural variations in the cavefish prechordal plate may relate to differential physical and adhesion properties of the organizer/prechordal cells in the two morphs. Live imaging will be necessary to better compare the properties of prechordal plate cells in cavefish and surface fish. Moreover, detailed analyses of expression of molecules involved in cell adhesion, such as snails and cadherins (Blanco et al., 2007; Montero et al., 2005; Shimizu et al., 2005), as well as those involved in membrane protrusion formation, such as β-actin (Giger & David, 2017), will help to explore the possibility that divergence in the intrinsic properties of prechordal plate cells may account for cavefish phenotypic evolution.

### Maternal control of gastrulation

Regardless of the striking morphological evolution observed in *A. mexicanus* morphotypes, their time of divergence has been estimated to be recent (less than 20.000 years ago) (Fumey et al., 2018). The inter-fertility, reminiscent of such a short divergence time between the two morphs, has allowed the use of hybrids for the identification of the genetic basis behind phenotypic change (Casane & Rétaux, 2016; Protas et al., 2006).

Since early embryonic development is driven by maternal determinants present in the oocyte before fecundation, the cross fertility in *A. mexicanus* species is a valuable tool to obtain information about the contribution of maternal effect genes to phenotypic evolution (Ma et al., 2018). Our analyses in F1 reciprocal hybrids demonstrate that the modifications in cavefish gastrulation are fully dependent on maternal factors. In line with this, our RNAseq analyses showed that the RNA composition of cavefish and surface fish eggs varied greatly, with 31.94% of the maternal genes expressed at the two-cell stage having differences in transcripts levels. Together these data strongly suggest that the eggs are a source of variation that can contribute to phenotypic evolution. In both RNAseq and qPCR analyses the candidate genes beta-catenin 1 and 2, involved in the establishment of the organizer (Kelly et al., 2000), did not show significantly different levels of expression. In contrast, two other genes, *oep* and *vsx1*, implicated in the development of the prechordal plate (Gritsman et al., 1999; Xu et al., 2014) showed reduced levels in cavefish compared to surface fish. However, overexpression of these two candidate genes by mRNA injection in cavefish was not able to recapitulate the gastrulation phenotype observed in the surface fish. Ma et al. (2018) have also recently described increased *pou2f1b*, *runx2b*, and *axin1* mRNA levels in unfertilized cavefish eggs as compared to surface fish eggs. These genes also show differential expression in our transcriptomic dataset. Classification of DEGs based on their biological role showed an enrichment on certain biological processes that may have been key for cavefish evolution. Relevant to this work we found that 6.5% of the “top DEGs with fold-change>5” are involved in signaling (Figure 8C). Some of these genes are regulators of the Wnt pathway (*i.e.* sFRP2, dkk2 and wnt11) that is important for the establishment of the embryonic organizer. Members of other signaling pathways are also greatly modified (i.e. FGF, BMP, Nodal). Our interpretation is that the origin of the induction of organizers with different properties in the two morphs might stem in an upstream maternally-regulated event, with a domino effect leading to morphological and functionally diverse brains. Our results on the impact of maternal determinants in forebrain morphogenesis are puzzling. Regarding the eye phenotype, our results are consistent with those of Ma et al., 2018 who examined maternal genetic effects in cavefish eye development and degeneration: at embryonic stages (12-24hpf), eye size and shape do not seem to be influenced by maternal factors. However, later larval lens apoptosis and eye regression appear to be under maternal control (Ma et al., 2018). This fits with our finding that lens size at 24hpf differs between reciprocal F1 hybrids. Indeed, the defective and apoptotic lens in cavefish is the trigger for eye degeneration (Yamamoto & Jeffery, 2000) and midline shh signaling indirectly impacts this lens-directed process (Hinaux et al., 2016; Ren et al., 2018; Yamamoto et al., 2004). Hence, the lens phenotype, but not the retina (which is in fact relatively properly formed and healthy in cavefish embryos), probably results from maternally-controlled developmental evolution in cavefish. This renders even more mysterious the long-standing question of what regulates the lens defects and apoptotic process in cavefish embryos.

More generally, regarding hypothalamic, eye and olfactory development that we have analyzed, our interpretation is that although maternal factors greatly influence early developmental decisions, later phenotypes become “diluted” due to other mechanisms entering into play. We suggest that once zygotic genome takes control over development, allelic dominance has a major impact on the phenotypes after 15 hpf onwards, although we could still observe some tendency to maternally-controlled phenotypes in hybrids for some of the traits analyzed. Of note in *Astyanax*, some behavioral traits in adults have already been shown to be under parental inheritance (Yoshizawa et al., 2012): the vibration attraction behavior and its underlying sensory receptors (the neuromasts) are under paternal inheritance in cavefish originating from the Pachón cave, while they are under maternal inheritance in cavefish originating from the Los Sabinos cave. These examples underscore the different levels of developmental regulation that must interact to produce a hybrid phenotype.

The study of the impact of maternal components in the morphological and developmental evolution of species is open. To our knowledge, besides *Astyanax* cavefish, only one study reported a maternal contribution regulating the developmental trajectory of entry into diapause in a killifish (Romney & Podrabsky, 2017). Thus *Astyanax* cavefish appear as a proper model to disentangle the very early genetic and embryonic mechanisms of morphological evolution. In addition, modified expression of maternal genes could be due to differential *cis*-regulation, which for maternal effect genes in the evolutionary context has not been explored yet to our knowledge in any species.

## Acknowledgements

We thank Stéphane Père, Victor Simon and Krystel Saroul for care of our *Astyanax* colony, and François Agnès and all other members of the DECA team for fruitful discussions and important suggestions to our study. This work has benefited from the facilities and expertise of the QPCR and sequencing platforms of I2BC, Gif sur Yvette. We thank Yan Jaszczyszyn from the I2BC sequencing platform for fruitful interactions. Grant support: FRM (Equipe FRM DEQ20150331745 RETAUX), CNRS and Becas Chile.

## Materials and methods

### A. mexicanus embryos

Our surface fish colony originates from rivers in Texas, United States, and our cavefish colony derives from the Pachón cave in San Luis Potosi, Mexico. Embryos were obtained by *in vitro* fertilization and/or natural spawnings induced by changes in water temperature (Elipot et al., 2014). Development of *A. mexicanus* at 24°C is similar and synchronous for both morphotypes (Hinaux et al., 2011). For this study morphological aspects were taken as strict criteria to stage the embryos (number of cells, percentage of epiboly and number of somites). *In vitro* fertilizations were performed to generate reciprocal F1 hybrids by fecundating cavefish oocyte with surface fish sperm (HybCF), and surface fish oocyte with cavefish sperm (HybSF).

### Whole mount in situ hybridization (ISH)

ISH was carried out as previously described (Pottin et al., 2011). Digoxigenin- and Fluorescein-labeled riboprobes were prepared using PCR products as templates. Genes of interest were searched in an EST (Expressed sequence tag) library accessible in the laboratory. Clones in library (pCMV-SPORT6 vector): *chordin* (ARA0AAA23YC10), *dkk1b* (ARA0AAA18YA07EM1), *eya2* (ARA0AAA19YL19EM1), *floating-head* (ARA0ACA35YA23), *myoD* (ARA0AAA95YG16), *msgn1* (ARA0ACA49YF15), *no-tail* (ARA0ABA99YL22), *npy* (FO263072), and *vsx1* (ARA0AHA13YJ18). Others: *fgf8* (DQ822511), *lhx9* (EF175738), *shh* (AY661431), *dlx3b (*AY661432), *hcrt (XM_007287820.3)*, *lhx7* (XM_022678613) cDNAs were previously cloned. Cloned for this study: total RNA from *Astyanax* embryos of various stages (2-24 hpf) was reverse-transcribed using the iScript cDNA synthesis kit (Bio-Rad) and amplified using the following primers:

brachyury, FP: CACCGGTGGAAGTACGTGAA, RP: GGAGCCGTCGTATGGAGAAG;

*lefty1*: FP: ACCATGGCCTCGTGCCTC; RP: TCAGACCACCGAAATGTTGTCCAC

Full length cDNAs were cloned into the pCS2+ expression vector using the indicated restriction sites:

*dkk1b* (sites EcoRI and XhoI), FP: GGTGGTGAATTCACCATGTGGCCGGCGGCGCTCTCAGCCCTGACCTTC, RP: ACCACCCTCGAGTCAGTGTCTCTGGCAGGTATGG;

*vsx1* (sites XhoI and XbaI), FP: GGTGGTCTCGAGACCATGGAGAAGACACGCGCG, RP: ACCACCTCTAGATCAGTTCTCGTTCTCTGAATCGC;

*oep* (*tdgf1*) (sites BamHI and XbaI), FP: GGTGGTGGATCCACCATGAGGAGCTCAGTGTTCAGG, RP: ACCACCTCTAGATCAAAGCAGAAATGAAAGGAGGAG.

### mRNA injections

*In vitro* transcription was carried out from PCR products using the SP6 RNA polymerase (mMESSAGE mMACHINE) to generate full length capped mRNA. Dilutions of the mRNA to 150-200ng/µL were prepared in phenol red 0.05%. Embryos at the one cell stage were injected with 5–10 nL of working solutions using borosilicate glass pipettes (GC100F15, Harvard Apparatus LTD) pulled in a Narishige PN-30 Puller (Japan).

### LiCl treatments

Embryos were enzymatically dechorionated in 1 mg/mL Pronase solution dissolved in EM, then they were incubated in LiCl solutions, 0.1 or 0.2M prepared in EM water, during the indicated time window. After the treatment, embryos were washed 5 times in EM, and we let them to develop until the desired stage for further analyses.

### Image acquisition and analyses

Whole mounted embryos stained by colorimetric and fluorescent ISH were imaged on a Nikon AZ100 multizoom macroscope coupled to a Nikon digital sight DS-Ri1 camera, using the NIS software. Mounted specimens were image on a Nikon Eclipse E800 microscope equipped with a Nikon DXM 1200 camera running under Nikon ACT-1 software. Confocal images were captured on a Leica SP8 microscope with the Leica Application Suite software. Morphometric analyses and cell counting were performed on the Fiji software (Image J). To measure the approximate extent of migration in the vegetal to animal axis (Height) we measured the distance from the margin to the leading cell normalized by the distance from the margin to the animal end (Figure 2 - figure supplement). To estimate the extent of dorsal convergence we measured either the width of the expression domain or the width of gap without expression (for example for *myoD*) normalized by the total width of the embryo (a representation using the expression of *myoD* at 70% epiboly is shown in Figure 3 - figure supplement). All measurements were normalized, unless otherwise indicated. Another means we used to calculate the width of expression was by measuring the angle (α) of the expression pattern from an animal view, using the center of the opposite site to the expression domain to set the vertex (a representation using the expression of *chordin* at 50% epiboly is shown in Figure 1 - figure supplement 1). To assess the width of the *pax2a* expression domain in the optic stalk/optic fissure we measured the angle (α) with the vertex set in the center of the lens (Figure 5 - figure supplement D) Statistical analyses were done in Graph pad prism 5.

### mRNA isolation

RNA pools were isolated from cavefish, surface fish and F1 hybrid embryos at the 2-cell stage (three independent biological replicates for each condition). Each sample corresponded to at least 20 embryos coming from two female individuals (40 embryos in total) to reduce inter-individual variability. Total RNA was extracted using TRIzol (Invitrogen, 2µL per embryo) and chloroform (0,2µL per µL of TRIzol) and purified with isopropanol (0,5µL per µL of TRIzol) and 70% ethanol and treated with DNase. Following purification, all samples were immediately quantified and assessed for RNA quality (A260/280 ratio ∼1.9– 2.1) using a NanoVue Spectrophotometer and stored at −80°C until use.

### qPCR

1 µg of total RNA was reverse transcribed in a 20 µL final reaction volume using the High Capacity cDNA Reverse Transcription Kit (Life Technologies) with RNase inhibitor and random primers following the manufacturer’s instructions. Quantitative PCR was performed on a QuantStudioTM 12K Flex Real-Time PCR System with a SYBR green detection protocol at the qPCR platform of the Gif CNRS campus. 3 µg of cDNA were mixed with Fast SYBRV R Green Master Mix and 500 nM of each primer in a final volume of 10 µL. The reaction mixture was submitted to 40 cycles of PCR (95°C/20 sec; [95°C/1 sec; 60°C/20 sec] X40) followed by a fusion cycle to analyze the melting curve of the PCR products. Negative controls without the reverse transcriptase were introduced to verify the absence of genomic DNA contaminants. Primers were designed using the Primer-Blast tool from NCBI and the Primer Express 3.0 software (Life Technologies). Primers were defined either in one exon and one exon–exon junction or in two exons span by a large intron. Specificity and the absence of multilocus matching at the primer site were verified by BLAST analysis. The amplification efficiencies of primers were generated using the slopes of standard curves obtained by a fourfold dilution series. Amplification specificity for each real-time PCR reaction was confirmed by analysis of the dissociation curves. Determined Ct values were then exploited for further analysis, with the *Gapdh* gene as reference.

### RNAseq analyses of maternal mRNAs

RNA sequencing was carried out at the I2BC High throughput sequencing platform (https://www.i2bc.paris-saclay.fr/spip.php?article399) using an Illumina NextSeq 500 sequencing instrument (version NS500446). All RNA samples were checked with a Bioanalyzer RNA 6000 pico chip (Agilent technologies) and passed quality threshold (RIN>9) prior to library preparation. Libraries were generated from purified total RNA using polyA selection (llumina TruSeq Stranded Protocol). Samples were sequenced for between 75 and 100 million reads (paired-end, 51-35bp) using the NextSeq 500/550 High Output Kit v2 (75 cycles). Following sequencing, raw data were retrieved (fastq-formatted files) and used for subsequent sequence alignment and expression analyses. Raw sequencing data are available through the NCBI Sequence Reads Archive (SRA) under BioProject accession PRJNA545230.

RNA-sequencing reads from each of the four conditions (surface fish, cavefish and reciprocal F1 hybrids) were trimmed using Cutadapt 1.15 and quality control was assessed using FastQC (v0.11.5). All downstream analyses were done using the European Galaxy Server (https://usegalaxy.eu, (Afgan et al., 2016)) with reverse (RF) strandness parameter. Read were aligned to the Surface Fish *Astyanax* genome (NCBI, GCA_000372685.2 Astyanax_mexicanus-2.0) using HISAT2 (Galaxy Version 2.1.0, (Kim et al., 2015)) and only perfectly aligned paired reads were kept for the following analysis (Filter SAM and Bam file Galaxy Version 1.8: Minimum MAPQ quality score 20 and Filter on bitwise flag “Read is paired” and “Read is mapped in a proper pair”). Then, aligned reads were counted using htseq-count (Galaxy Version 0.9.1, (Anders et al., 2015)) and the *Astyanax mexicanus* annotation from NCBI release 102 (https://www.ncbi.nlm.nih.gov/genome/?term=txid7994[orgn]). Differentially-expressed genes (DEG) between cavefish and surface fish were detected using DESeq2 (Galaxy Version 2.11.40.6, (Love et al., 2014)). Based on the PCA and sample to sample distance analyses, data from F1 hybrid were combined with their respective mother morphotype for the DEG analysis. Only genes with a FDR<0.01 (false discovery rate, p value adjusted for multiple testing with the Benjamini-Hochberg procedure) and absolute fold change higher than 1.5 (log2(FC)>0.58) were kept as significantly over- or under-expressed in cavefish compared to surface fish. Mapped reads were visualized using the genome browser IGV (http://www.broadinstitute.org/igv/) (Robinson et al., 2011).

A Gene Ontology Annotation file for the *Astyanax* transcriptome (from NCBI database) was generated using OmicsBox (formerly Blast2GO, https://www.biobam.com/) following the general workflow presented by the software: BLAST with CloudBlast (restricted to the teleosteii database, keeping the top 20 results with an e-value of 10^(-5)), followed by mapping (GO version April 2019), annotation and InterProScan analysis in parallel. The annotation file was generated by merging the annotated BLAST results with InterProScan results. Gene Ontology Enrichment analysis on differentially-expressed genes (FDR<0.01 and FC>1.5) was carried out on Galaxy using GOEnrichment (Galaxy Version 2.0.1) with p-value cut-off of 0.01. We used several thresholds of fold change (FC>1.5, FC>5, FC>10, FC>20 and FC>50) to define gene sets and performed the analysis using the genes expressed at 2-cell stage as reference (n=20730). In this study, on the study gene set with FC>5 was kept as it is the most biologically meaningful.

---

**Figure 1 - figure supplement 1.**
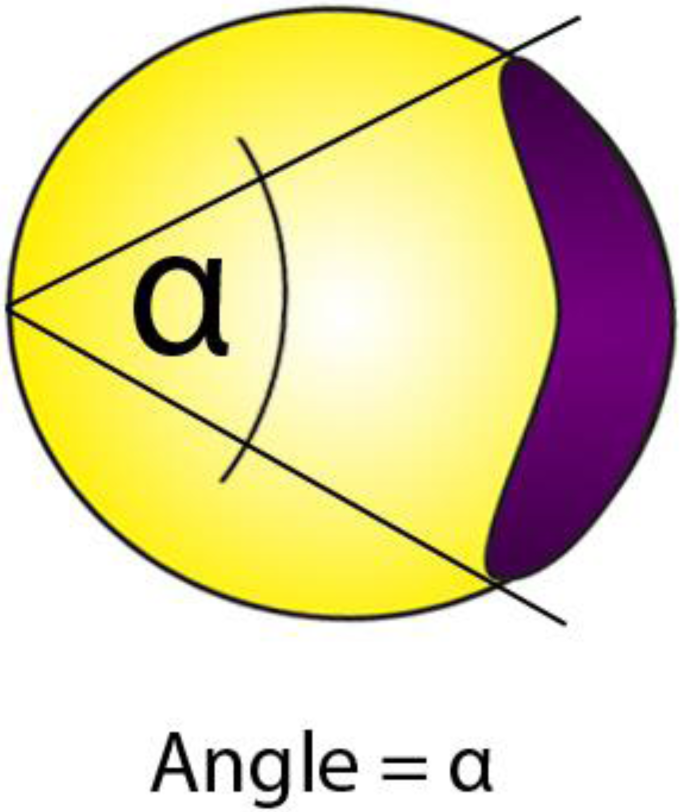
Measurement of angle in embryos stained for *chordin* at 50% of epiboly, in animal view.

**Figure 1 - figure supplement 2.**
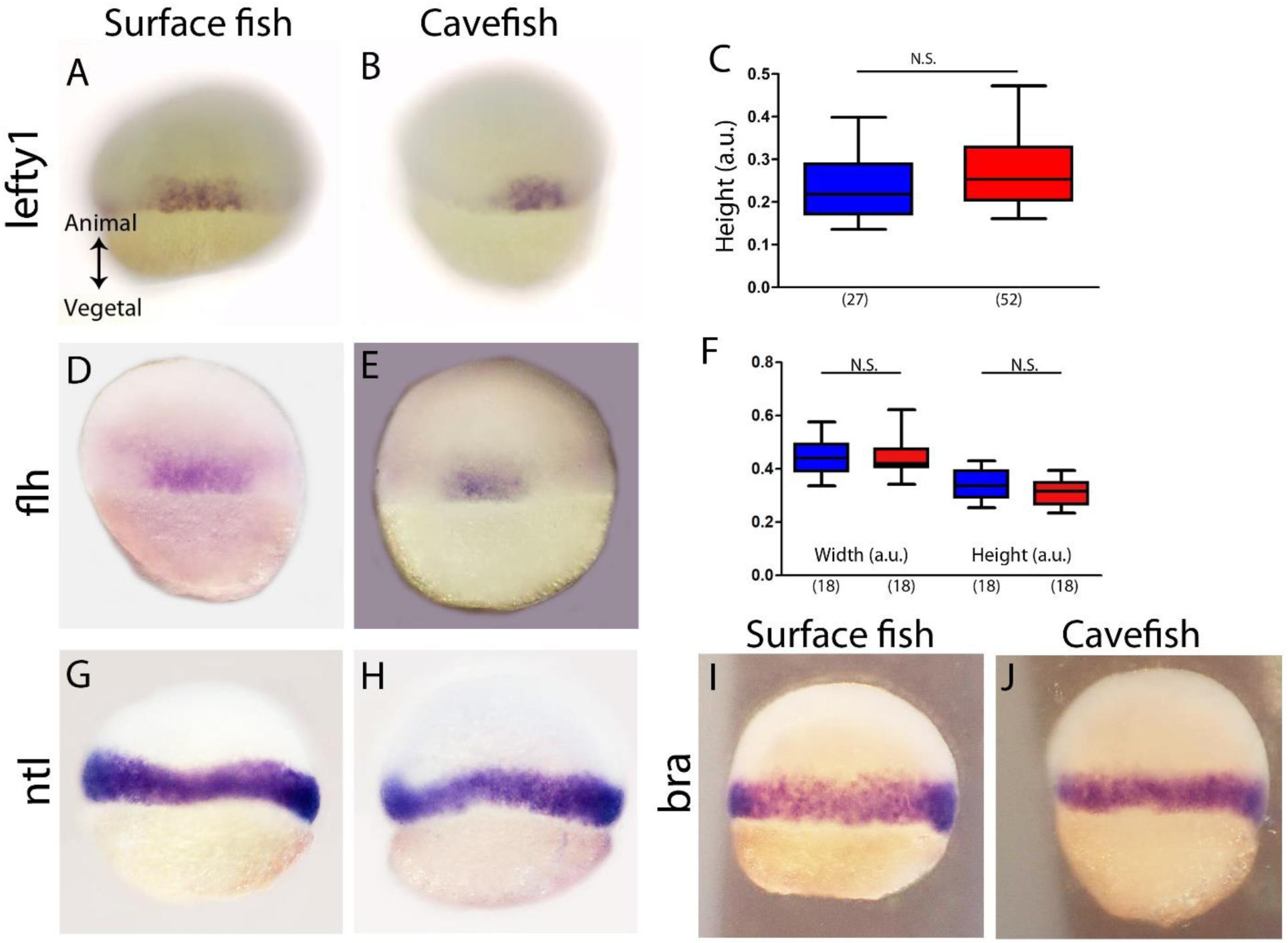
(A-C) Expression of *lefty1* in surface fish (A) and cavefish (B) in dorsal view. (C) Quantification of the height for *lefty1* expression. (D-F) Expression of *flh* in surface fish (D) and cavefish (E) in dorsal view. (F) Quantification of the width (left) and the height (right) of *flh* expression. (G-H) Expression of *ntl* in surface fish (G) and cavefish (H). (I-J) Expression of *bra* in surface fish (I) and cavefish (J). All embryos in dorsal view, animal pole upwards.

**Figure 2 - figure supplement.**
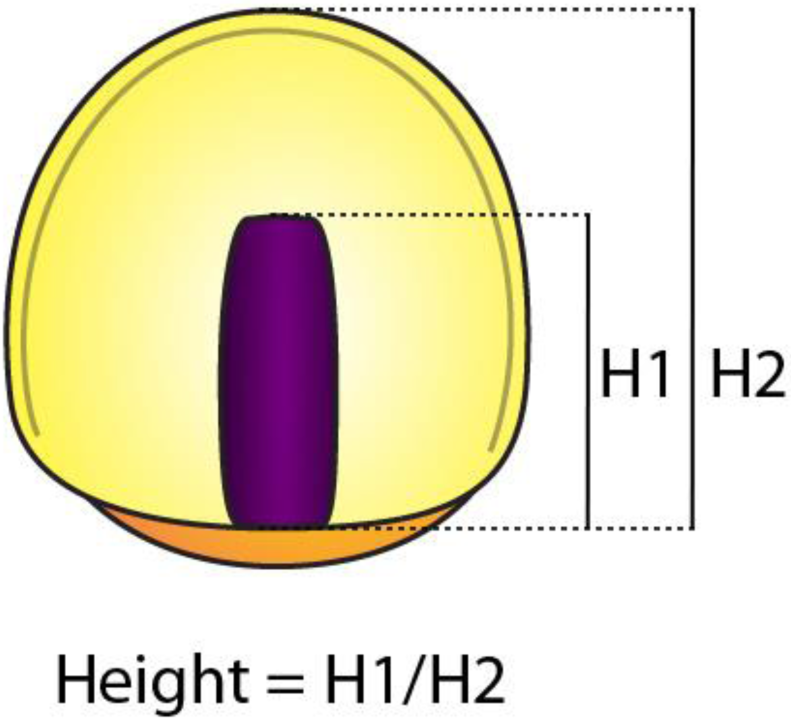
Measurement of the height, corresponding to the ratio between H1 (distance from the margin to the leading cell) and H2 (distance from the margin to the animal pole).

**Figure 3 - figure supplement.**
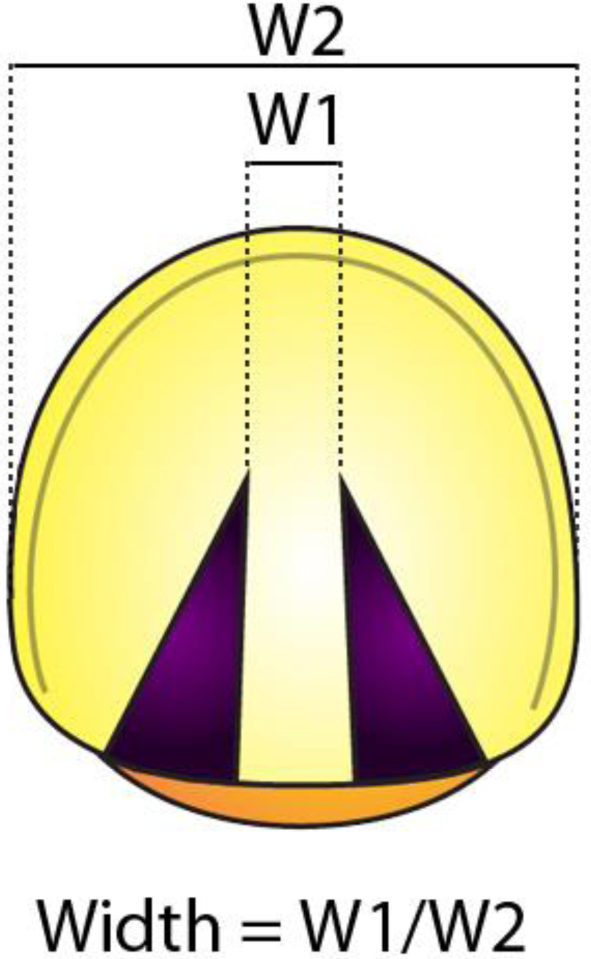
Measurement of the width, corresponding to the ratio between W1 (distance of the central gap in this case, or the expression domain) and W2 (the total width of the embryo)

**Figure 4 - figure supplement 1.**
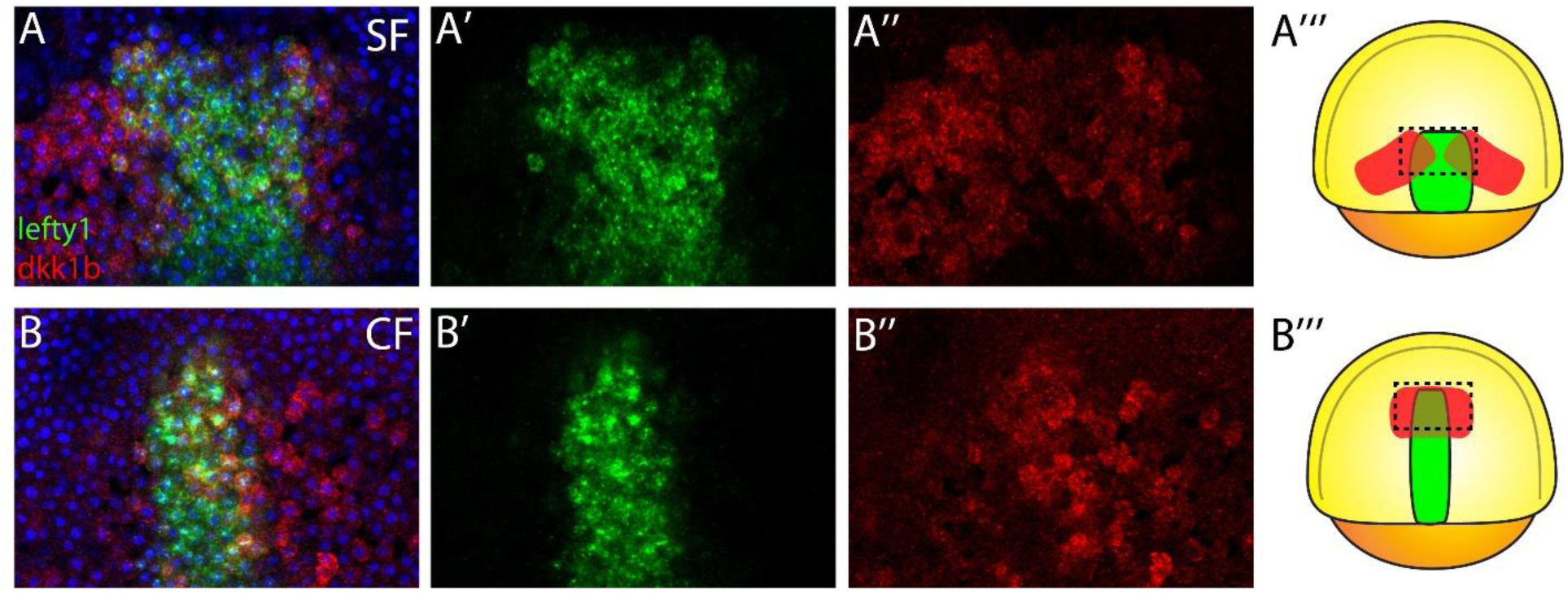
Expression of *dkk1b* (red) and *lefty1* (green) in surface fish (A) and cavefish (B) at 70% of epiboly. Scheme of surface fish (A’’’) and cavefish (B’’’) at 70% of epiboly with the region of interest indicated in dashed line.

**Figure 4 - figure supplement 2.**
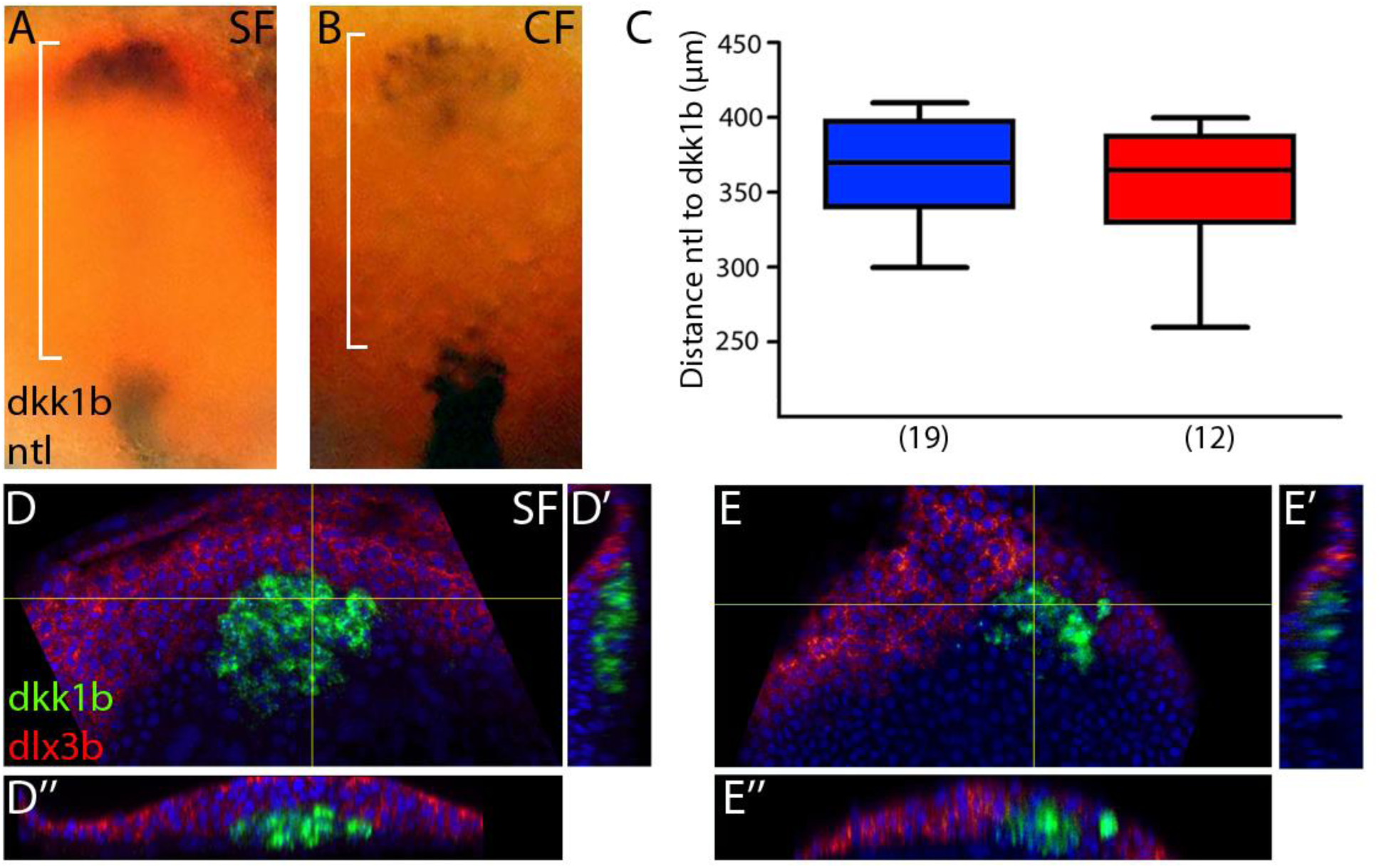
(A-C) Expression of *dkk1b* and *ntl* (notochord) at 10hpf in surface fish (A) and cavefish (B). Quantification of the distance from the leading notochordal cell to the leading polster cell (C) in a dorsal view, indicated in brackets in A and B. (D-E) Confocal images of the expression of *dkk1b* (green) and *dlx3b* (red, neural plate border) at 10hpf in surface fish (D) and cavefish (E). D and E are projections of 3µm, D’ and E’ are reconstructions of sagittal section (yellow line, vertical), and D’’ and E’’ are reconstructions from a transverse section (yellow line, horizontal). A and B are whole mounted embryos, and D and E are dissected embryos.

**Figure 5 - figure supplement.**
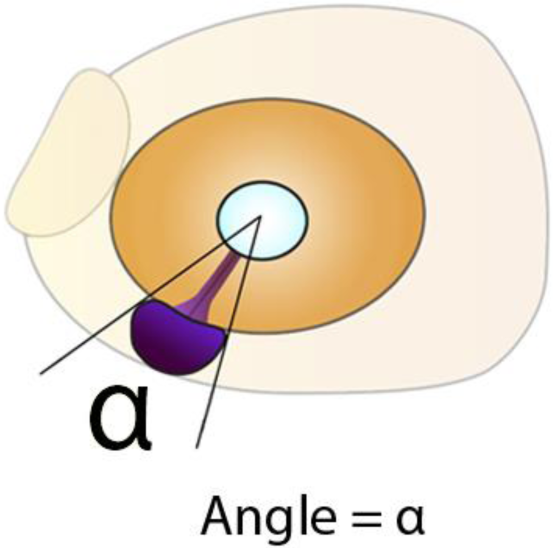
Measurement of angle in embryos stained for *pax2a* at 36hpf, in lateral view.

**Figure 6 - figure supplement.**
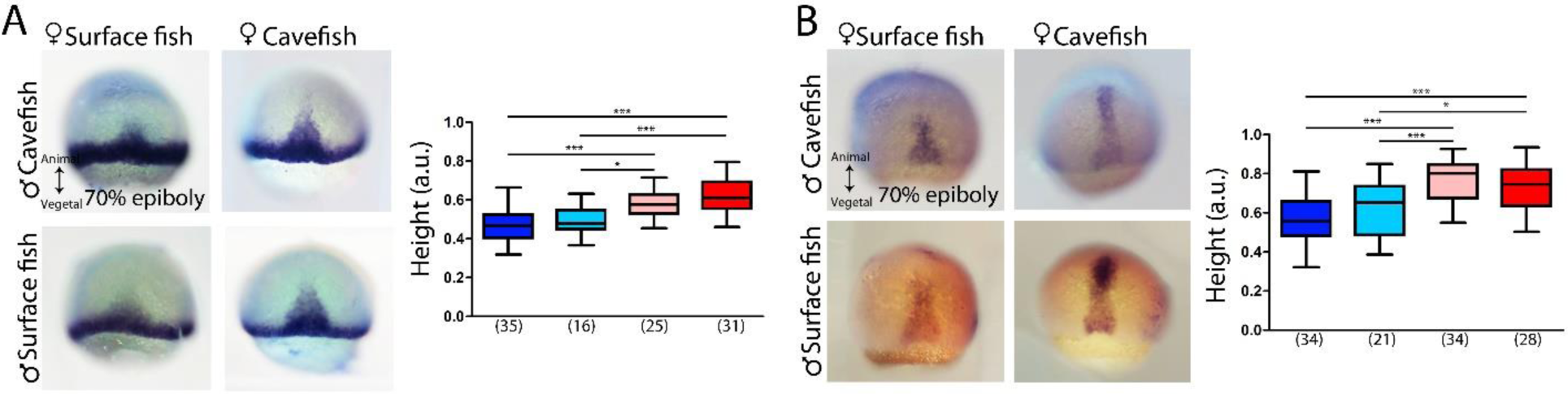
Expression of *ntl* (A) and *lefty1* (B) at 70% of epiboly (B). In the panels A-B are shown HybSF (top left), cavefish (top right), surface fish (bottom left) and HybCF (bottom right). Quantification of height in *ntl* (A, right) and *lefty1* (B, right) labeled embryos at 70% epiboly. Color code: surface fish, blue; HybSF, light blue; HybCF, pink; and cavefish, red.

**Figure 8 - figure supplement.**
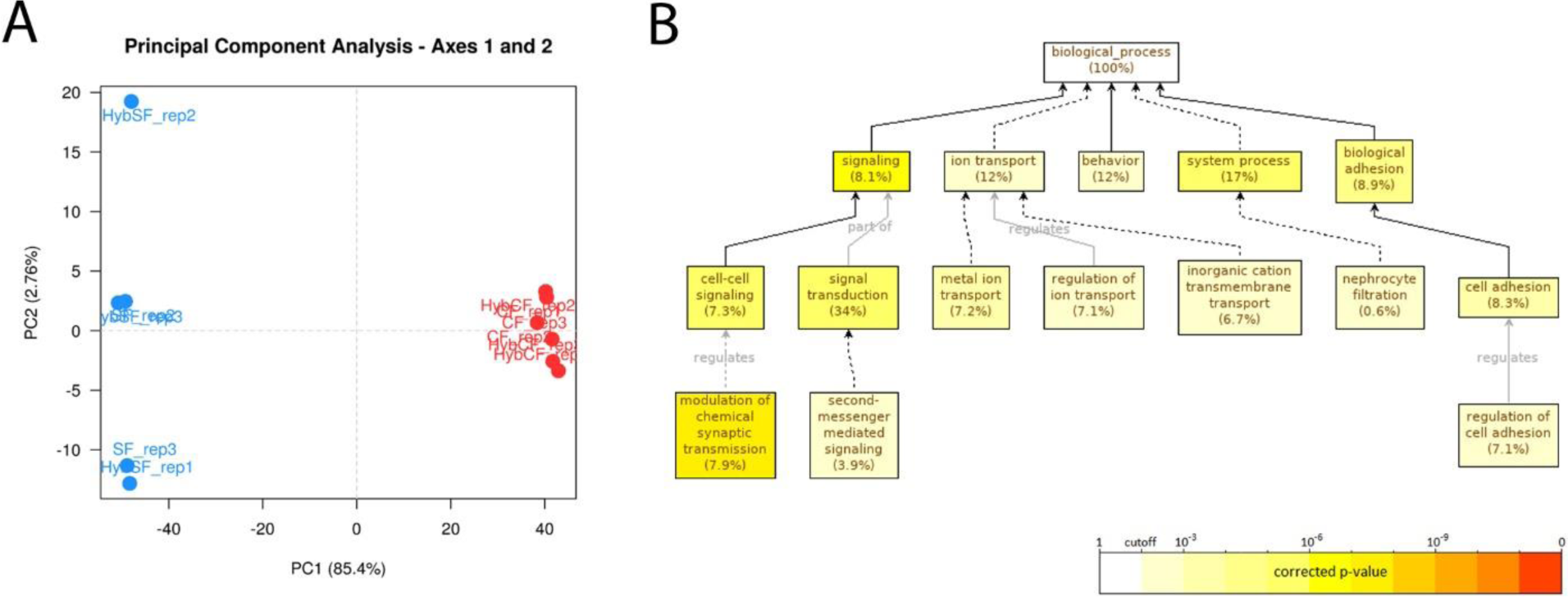
(A) Principal Component Analysis (PCA) of all the samples for PC1 and PC2. Blue dots correspond to samples coming from a female surface fish and red dots samples coming from female cavefish irrespective of the male morphotype. Note that PC1 and PC2 represent 85.4% and 2.7% of the variation, respectively. (B) Go Enrichment graph for a subset of cavefish up-regulated DEGs, with a fold change higher than 5. Black lines correspond to “is_a” relationship whereas grey lines correspond to the annotated relationship. Full lines correspond to direct relationship and dashed line to indirect relationship (i.e. some nodes are hidden). The color of a node refers to the adjusted p-value (FDR) of the enriched GO term and the percentage corresponds to the frequency of the GO term in the studied gene set at the level considered. A given gene can have several GO terms. Only enriched GO terms that pass the threshold (p-value<0.01) are displayed on the graph.

## Notes

#### Summary of Updates

This version of the manuscript has been revised to update the dataset: 1) functional experiments manipulating Wnt signaling to probe the role of the described dkk1b hetechrony 2) RNA-seq on 2-cell stage embryos (surface fish, cavefish and reciprocal F1 hybrids) to demonstrate molecularly the strong maternal contribution.

